# Combined loss of obscurin and obscurin-like 1 in murine hearts results in diastolic dysfunction, altered metabolism and deregulated mitophagy

**DOI:** 10.1101/2022.08.24.505098

**Authors:** Kyohei Fujita, Patrick Desmond, Jordan Blondelle, Matúš Soták, Meenu Rohini Rajan, Madison Clark, Eric Esteve, Yunghang Chan, Yusu Gu, Valeria Marrocco, Nancy D. Dalton, Majid Ghassemian, Aryanne Do, Matthew Klos, Kirk L. Peterson, Farah Sheikh, Yoshitake Cho, Emma Börgeson, Stephan Lange

**Author notes:** Correspondence should be directed to Dr. Stephan Lange.

## Abstract

Muscle proteins of the obscurin protein family play important roles in sarcomere organization, sarcoplasmic reticulum (SR) and T-tubule architecture and function. However, their precise molecular functions and redundancies between protein family members as well as their involvement in cardiac diseases remain to be fully understood.

To investigate the functional roles of obscurin and its close homologue obscurin-like 1 (Obsl1) in the heart, we generated and analyzed knockout mice for obscurin, Obsl1, as well as obscurin/Obsl1 double-knockouts (dKO). We show that dKO mice are viable but show postnatal deficits in cardiac muscle SR and mitochondrial architecture and function at the microscopic, biochemical and cellular level. Altered SR structure resulted in perturbed calcium cycling, while mitochondrial ultrastructure deficits were linked to decreased levels of Chchd3, a Micos complex protein. Hearts of dKO mice also show increased expression of Atg4d, a novel Obsl1 interacting protein, resulting in abnormal mitophagy and increased unfolded protein response. At the physiological level, loss of obscurin and Obsl1 resulted in a profound delay of cardiac relaxation, associated with metabolic signs of heart failure.

Taken together, our data suggest that obscurin and Obsl1 play crucial roles in cardiac SR structure, calcium cycling, mitochondrial function, turnover and metabolism.

## Introduction

Giant muscle proteins have long been known to serve a multitude of roles for the organization and function of cross-striated muscles. The best investigated example is the giant protein titin, which, among its other functions, acts as a blueprint needed for sarcomerogenesis, a tunable passive spring element, and a docking-hub for a multitude of accessory proteins to maintain sarcomere integrity and enable muscle-specific signaling^1–6^. A family of proteins with similar domain layouts and comparably ‘outsized’ roles in the organization and function of muscles is the obscurin protein family^7–9^.

Like titin, obscurin, the initial member of this protein family, consists of serially linked immunoglobulin and fibronectin domains in addition to C-terminally located signaling domains ^7–10^. The protein exists in at least three major splice isoforms (obscurin-A, obscurin-B and obscurin associated kinase KIAA1639) and shows a high degree of evolutionary conservation, as unc-89, the invertebrate homologue of obscurin, has comparable roles for muscle formation and function as its vertebrate counterpart^11–13^.

Obscurin serves multiple functions in cross-striated muscles. It promotes the integrity of the sarcomere by interacting with several structural proteins, perhaps also reflected in its varied localization along myofibers. Obscurin localizes initially to Z-disks in embryonic hearts, but later is mostly found at the sarcomeric M-band^7^. The Z-disc localization is mediated by interaction of obscurin domains Ig58-59 with titin Ig9^14^, while its presence at M-bands is the result of obscurin Ig1 association to titin Ig-domain M10^15^. Mutations in titin Ig-domain M10 that weaken binding of obscurin (and Obsl1) are linked to the development of limb-girdle muscular dystrophy (LGMD2J)^15^. Obscurin also interacts with Rho-A, tying it to the M-band^16, 17^. While this interaction may play a role in mechanosensing of non-muscle cells^18^, cardiac roles for obscurin binding to Rho-A remain to be better understood. Giant obscurin isoforms also link sarcomeres to membranes, by interacting at their N-terminus with titin and myomesin^15, 19^, while the unstructured C-terminus in obscurin-A binds to ankyrin isoforms at the sarcoplasmic reticulum (SR) or sarcolemma membranes^17, 20–25^. Rho-GEF domains in obscurin-A and -B as well as several smaller novel obscurin isoforms were also shown to interact with phospholipid components of membranes^26^. Together, these interactions suggest important roles for the organization of membrane and membrane-associated proteins as well as the architecture of the SR and sarcolemma, indirectly also modulating calcium cycling and ion homeostasis.

Obscurin knockouts develop a mild myopathy characterized by increased numbers of centralized nuclei, but no discernable cardiac phenotype^23^. However, several mutations in human obscurin are linked to the development of multiple forms of cardiac and skeletal muscle myopathies in patients^27–31^. One of the best characterized gene variants in obscurin that is associated with hypertrophic cardiomyopathy in patients is the R4344Q mutation^32^. Specifically, this mutation in obscurin’s Z-disk interacting domains Ig58-59 has been shown to influence cardiac calcium cycling by modulating binding to the sarcoplasmic/endoplasmic reticulum calcium ATPase 2 (Serca2) associated micropeptide phospholamban (Pln)^33^, although the relevance of this interaction is contested^34^. Deletion of obscurin Ig58-59 resulted in an arrhythmogenic cardiomyopathy in aged mice^33^. Patients with bi-allelic loss of obscurin were recently shown to predispose affected individuals to severe recurrent rhabdomyolysis^35^. Consequently, these patients are exercise intolerant. However no cardiac phenotype was noted in affected individuals as judged by echocardiography and MRI, suggesting that loss of obscurin may not represent a substantial cause of cardiomyopathies.

Obscurin-like 1 (Obsl1) and striated muscle preferentially expressed gene (also known as striated muscle enriched protein kinase, Speg) are the two other vertebrate members of the obscurin family, having been evolutionarily formed by duplication of the 5’ and 3’ ends of the obscurin gene sequence, respectively^11^.

Functions for Speg in cardiac and skeletal muscles have been intensively investigated. Its main role lies in the organization and phosphorylation of junctional SR proteins as well as regulation of muscle cell calcium homeostasis. Loss of Speg results in a severe dilated cardiomyopathy phenotype in mice ^36^. Patients with pathological mutations in Speg manifest with symptoms that can range from dilated cardiomyopathy^37–40^ to centronuclear myopathy^39, 41, 42^. Like obscurin-B, Speg also harbors a set of tandem protein kinases that are catalytically active, and whose function is partially conserved throughout evolution^43–46^. Identified substates for tandem kinases in obscurin and Speg include the ryanodine receptor, junctophilin, the phosphatase myotubularin (Mtm1) or Serca2 ^39, 47–49^. Speg kinase domains are able to autophosphorylate the protein^50^, similar to obscurin-B kinase domain 1, which was also shown to autophosphorylate a low complexity region located between the tandem kinases^51^. The inter-kinase region in unc-89 was recently shown to act as entropic spring element in nematodes^52^, suggesting similar functions for the vertebrate Speg and obscurin. Moreover, both Speg and obscurin are heavily modified during maximal intensity contractions and in exercise, including in the inter-kinase region^53, 54^.

Comparatively little is known for Obsl1 functions in cross-striated muscles. Obsl1 was first reported as a cytoskeletal adapter protein with various subcellular localizations in cross-striated muscle cells^15, 43, 55^. Similar to obscurin, Obsl1 binds in its N-terminus also to titin and myomesin, providing functional redundancy for these interactions^15, 19^. However, it seems to lack binding sites to ankyrin or other proteins associated with the SR or sarcolemma membrane, offering instead binding sites to myosin, cytoskeletal or accessory proteins, such as filamin C or dystonin^43, 56^. Global loss of Obsl1 in mice results in early embryonic lethality before E17, underscoring important functions of the protein outside of muscle tissues^43^. Pathological variants of Obsl1 in humans are associated with 3M syndrome, a growth disorder that is also linked to mutations in ccdc8 and cullin-7^57–59^.

Little is known how loss of obscurin and/or Obsl1 would affect cardiac development and function. Data from patients with bi-allelic loss of obscurin or patients suffering from 3M syndrome caused by mutations in Obsl1 had unremarkable cardiac phenotypes. In this study, we set out to study the phenotype of obscurin and Obsl1 cardiac-specific single and double knockout (dKO) mice. Our investigations uncovered that loss of either obscurin or Obsl1 did not alter viability of mice. However, the combined cardiac-specific loss of obscurin and Obsl1 in dKO mice led to postnatal cardiac defects including severe diastolic dysfunction and heart failure associated with premature death. Our molecular and biochemical analyses uncovered changes to SR structure and calcium cycling, which could be mostly tied to loss of obscurin. Conversely, dKO mice showed additional changes to cardiac metabolism, mitochondrial structure and metabolism as well as turnover. Mitochondrial defects are mechanistically driven by loss of interactions between Obsl1 and the mitochondrial Micos complex protein Chchd3 as well as Obsl1 and the autophagy related and mitochondrially targeted Atg4d cysteine peptidase. As a result of the mitochondrial stress, dKO hearts display increased expression of proteins involved in the unfolded protein response (UPR).

## Results

### Obsl1 and/or obscurin are not required for cardiac sarcomerogenesis and development

Global Obsl1 knockout mice are embryonically lethal^43^, making them unsuitable to investigate cardiac-specific roles for Obsl1 and test for functional redundancy between obscurin and Obsl1. To overcome the lethality, we crossed conditional Obsl1 mice with Cre recombinase expressing mice under control of a transgenic Nkx2.5 promoter^60^. These mice develop normally and display no overt signs of cardiomyopathy or premature death (**Figures 1A, 1B**). A similar finding was observed in obscurin knockout mice, which also lack an apparent cardiac phenotype^23^. To investigate if the lack of a cardiac phenotype might be due to a rescue by the other obscurin protein family member, we generated cardiac-specific obscurin/Obsl1 double knockouts (dKO; **Figure 1C**). Surprisingly, dKO mice developed normally without any overt phenotypical defects at birth. However, starting from 6 months of age, dKO mice show a gradual worsening in survival (**Figure 1A**). Hearts of dKO did not display clear morphological indications for cardiac pathology (e.g., hypertrophy or dilatation) and no molecular indication for defects to the structural proteins of the sarcomeric Z-disc and M-band were observed (**Figures 1B, 1D**). While loss of Obsl1 alone had no effect on obscurin localization to sarcomeric M-bands (**Figure S1A**), its protein levels were increased (**Figure 1C**). However, loss of obscurin alone did result in a shift of Obsl1 from the M-band to both, M-band and Z-discs (**Figure S1B**), without affecting Obsl1 protein levels (**Figure 1C**). Speg, the third member of the obscurin protein family, was found to localize to the region of the Z-disc in control hearts, and shifted its subcellular localization to Z-discs and intercalated discs in either obscurin knockout, Obsl1 knockout, or dKO hearts (**Figure S1C**). In addition, SPEG protein levels increased with loss of Obsl1 (**Figure 1C**). We also noted a reduction in the EH-isoform of myomesin-1, which is thought to be biomechanically compliant due to the EH-domain acting as an entropic spring element^61–63^. However, overall protein levels of the Obsl1/obscurin binding partner myomesin-1 remained unchanged (**Figure S1D**).

**Figure 1.**
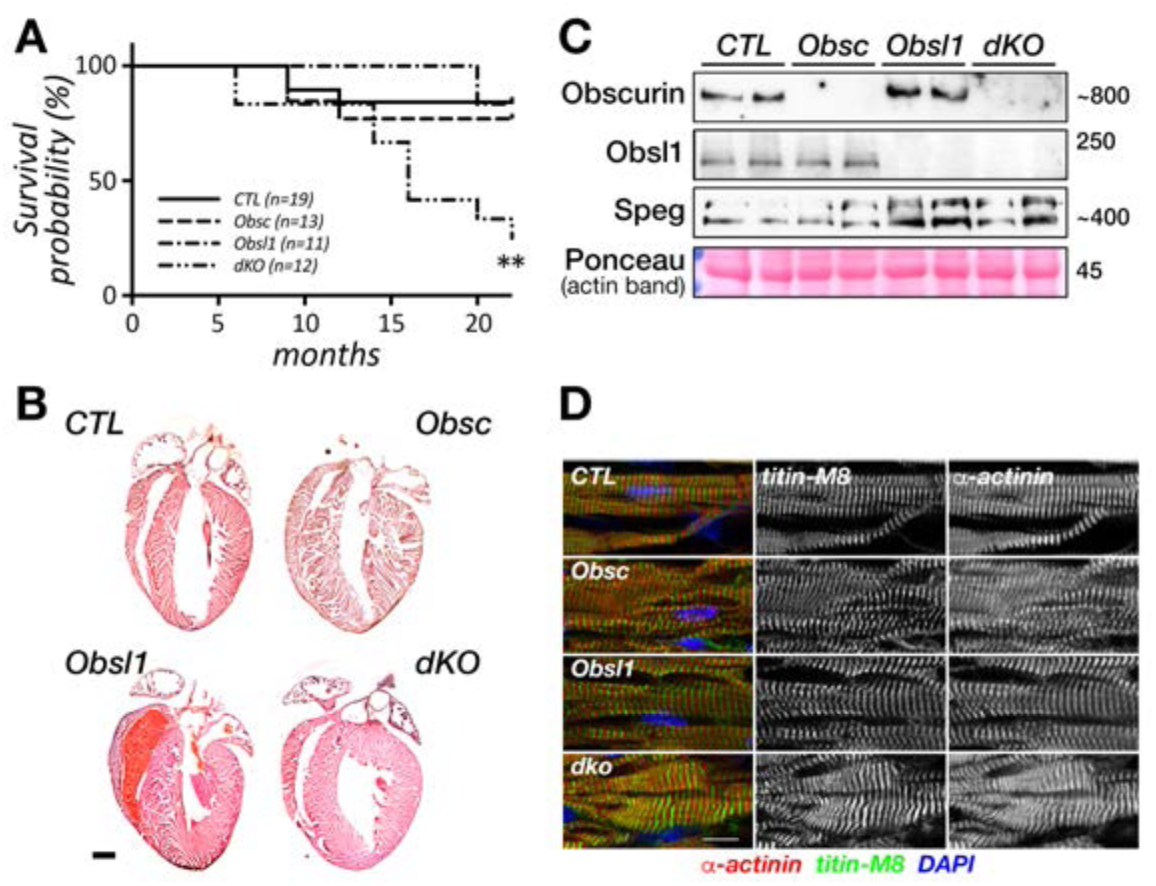
Cardiac ablation of obscurin (Obsc) and Obsl1 is associated with reduced life expectancy in mice. A. Kaplan-Meier survival probability curve for cardiac-specific obscurin (Obsc), Obsl1 and obscurin/Obsl1 double knockouts (dKO) compared to wildtype controls (CTL). ** p < 0.01 using Mantel-Cox test. B. Hematoxylin-Eosin stained sections of hearts from cardiac-specific obscurin, Obsl1 and obscurin/Obsl1 double knockouts (dKO) compared to wildtype controls (CTL). Scalebar = 1mm. C. Immunoblot analysis of cardiac extracts from cardiac-specific obscurin, Obsl1, obscurin/Obsl1 double knockouts (dKO) and wildtype controls (CTL) probed with antibodies against obscurin, Obsl1 and Speg. Ponceau stained actin is the loading control. D. Immunofluorescence of frozen cardiac sections from cardiac-specific obscurin, Obsl1, obscurin/Obsl1 double knockouts (dKO) and wildtype controls (CTL) decorated with antibodies against sarcomeric ⍺-actinin-2 (red) and titin-M8 (green), marking the sarcomeric Z-disc and M-band, respectively. DAPI (blue) was used as a counterstain. Scalebar = 20µm.

### Loss of obscurin/Obsl1 results in diastolic dysfunction

To determine the cause of premature death in dKO mice, we employed transthoracic echocardiography of male and female control (wildtype), obscurin knockout, Obsl1 knockout and dKO animals (**Figure 2, Table 1**). Cardiac functions as assessed by % fractional shortening were comparable between wildtype and all knockouts at various ages (**Figure 2A, Table 1**). Chamber dimensions (left ventricular internal dimensions in diastole and systole; LVIDd and LVIDs, respectively) and wall thicknesses (interventricular septal and left ventricular posterior wall thicknesses in diastole; IVSd and LVPWd, respectively) also remained largely comparable to controls. We only found transient changes in IVSd and LVPWd thicknesses of obscurin and dKO hearts at 6-9 months, with dKO displaying a moderate hypertrophy, while obscurin knockouts show a reduction in wall thicknesses.

**Figure 2.**
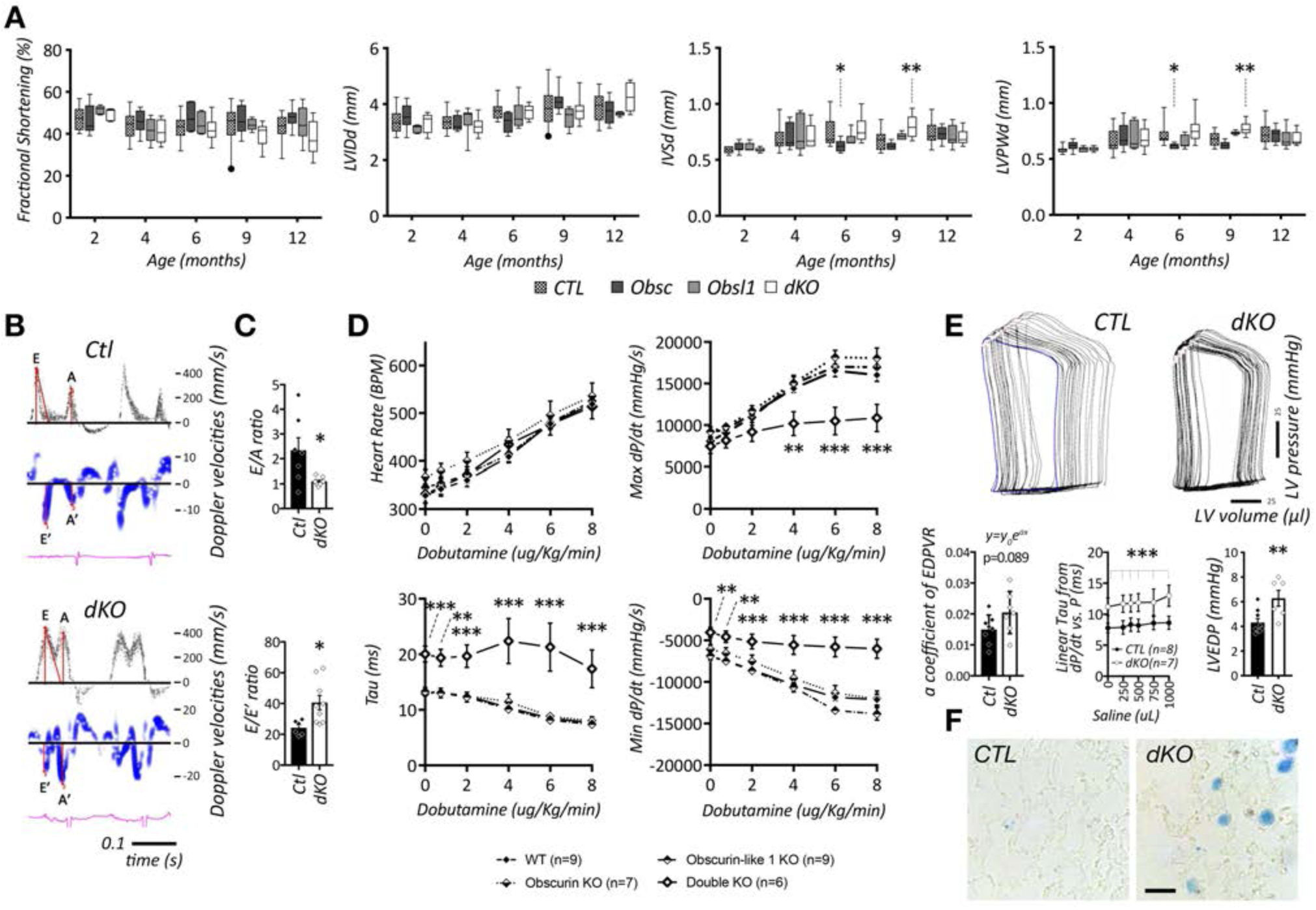
Cardiac physiology of cardiac-specific obscurin (Obsc), Obsl1 and obscurin/Obsl1 double knockouts (dKO) compared to wildtype controls (CTL). A. Fractional shortening (% FS), left ventricular internal diameter in diastole (LVIDd), interventricular septal thicknesses in diastole (IVSd), and left ventricular posterior wall thicknesses in diastole (LVPWd), of all groups at different ages as determined by transthoracic echocardiography. Box and whiskers plots depict median values with 5-95 percentile whiskers and outliers as individual data points. * p < 0.05, ** p < 0.01 vs. CTL by two-way ANOVA analyses, with multiple comparisons using Dunnett’s test. Sample sizes and additional parameters can be found in Table 1. B-C. Doppler and pulsed wave echocardiography of wildtype (CTL) and dKO mice to measure ratio of peak velocity blood flow from gravity in early diastole to peak velocity flow in late diastole (E/A ratio; C top panel), and ratio of mitral valve velocity of early filling to early diastolic mitral annular notion (E/E’ ratio; C bottom panel). Shown are averages with standard error of mean values. * p < 0.05 by T-test. Further data can be found in Table 2. D. Hemodynamics analyses of all groups at baseline and with increasing dobutamine amounts. Shown are changes to heart rate (HR in bpm), Max dP/dt, Min dP/dt (in mmHg/s) as well as Tau (in ms). Shown are averages with standard error of mean values. * p < 0.05, ** p < 0.01, *** p < 0.001 vs. CTL by two-way ANOVA analyses, with multiple comparisons using Dunnett’s test. Further data can be found in Table 3 and Supplemental Figure S2. E. Pressure volume (P/V) loops (E top panel) of wildtype (CTL) and dKO mice. Calculation of alpha-coefficient of fitted end-diastolic pressure volume relationship (EDPVR; E bottom left panel), linear *Tau* as measured by dP/dt vs. P (E bottom middle panel) and left ventricular and diastolic pressures (LVEDP; E bottom right panel). Shown are averages with standard error of mean values. ** p < 0.01, *** p < 0.001 vs. CTL by T-test. F. Prussian blue staining of wildtype (CTL) and dKO mouse lung sections. Scalebar = 20µm.

**Table 1.** Echocardiographic analysis of cardiac function in control (CTL), obscurin knockout (Obsc), Obsl1 knockout and dKO male and female mice. Shown are averages and standard deviation (SD) values. * p < 0.05, ** p < 0.01 by two-way ANOVA analyses, followed by Dunnett’s multiple comparisons test. Abbreviations: HR – heart rate, FS – fractional shortening, LVIDd – left ventricular internal dimensions in diastole, LVIDs – left ventricular internal dimensions in systole, IVSDd – interventricular septal thickness in diastole, LVPWd – left ventricular posterior wall thickness.

|  | Age (months) | Sample size | HR (bpm) | FS (%) | LVIDd (mm) | LVIDs (mm) | IVSd (mm) | LVPWd (mm) |
| --- | --- | --- | --- | --- | --- | --- | --- | --- |
| CTL | 2 | 15 | 552±98 | 47.7±5.5 | 3.4±0.5 | 1.8±0.32 | 0.59±0.02 | 0.59±0.03 |
|  | 4 | 19 | 593±75 | 44.0±6.4 | 3.4±0.3 | 1.9±0.26 | 0.68±0.11 | 0.66±0.11 |
|  | 6 | 17 | 601±73 | 43.3±5.5 | 3.8±0.3 | 2.1±0.26 | 0.75±0.12 | 0.71±0.09 |
|  | 9 | 19 | 541±85 | 44.6±8.4 | 3.8±0.6 | 2.1±0.58 | 0.68±0.08 | 0.67±0.06 |
|  | 12 | 18 | 560±54 | 42.8±6.5 | 3.9±0.5 | 2.2±0.49 | 0.75±0.10 | 0.72±0.08 |
| Obsc | 2 | 11 | 476±89 | 46.5±6.6 | 3.6±0.5 | 1.9±0.39 | 0.61±0.05 | 0.62±0.05 |
|  | 4 | 16 | 526±70* | 45.9±5.3 | 3.3±0.3 | 1.8±0.20 | 0.72±0.11 | 0.70±0.11 |
|  | 6 | 5 | 511±42* | 48.0±7.1 | 3.4±0.4 | 1.8±0.35 | 0.62±0.06* | 0.61±0.03 |
|  | 9 | 7 | 478±67 | 46.7±7.4 | 4.1±0.4 | 2.2±0.49 | 0.62±0.03 | 0.63±0.03 |
|  | 12 | 11 | 530±56 | 47.5±4.3 | 3.7±0.5 | 2.0±0.35 | 0.73±0.07 | 0.71±0.06 |
| Obsl1 | 2 | 5 | 662±57 | 51.4±2.0 | 3.1±0.2 | 1.5±0.02 | 0.61±0.04 | 0.58±0.03 |
|  | 4 | 8 | 608±78 | 41.4±5.5 | 3.4±0.5 | 2.0±0.41 | 0.73±0.16 | 0.71±0.13 |
|  | 6 | 6 | 572±62 | 44.9±5.4 | 3.6±0.5 | 2.0±0.46 | 0.68±0.06 | 0.65±0.06 |
|  | 9 | 5 | 602±32 | 44.5±2.8 | 3.5±0.4 | 2.0±0.23 | 0.71±0.03 | 0.73±0.02 |
|  | 12 | 7 | 594±30 | 45.0±8.4 | 3.6±0.1 | 2.0±0.32 | 0.70±0.10 | 0.68±0.09 |
| dKO | 2 | 5 | 541±39 | 49.4±3.0 | 3.3±0.4 | 1.7±0.16 | 0.59±0.02 | 0.59±0.02 |
|  | 4 | 10 | 542±56 | 41.0±5.4 | 3.2±0.3 | 1.9±0.20 | 0.72±0.11 | 0.68±0.10 |
|  | 6 | 10 | 568±50 | 42.0±5.6 | 3.7±0.2 | 2.2±0.29 | 0.77±0.12 | 0.76±0.12 |
|  | 9 | 13 | 530±44 | 39.7±5.8 | 3.8±0.4 | 2.3±0.34 | 0.80±0.10** | 0.77±0.06** |
|  | 12 | 5 | 519±57 | 38.3±8.7 | 4.3±0.5 | 2.7±0.66 | 0.70±0.08 | 0.68±0.07 |

To gain further insight into the cardiac physiology, we performed pulsed wave and tissue Doppler echocardiography^64^, measuring the E-wave, which represents early passive filling of the left ventricle (LV), and the A-wave, representing active filling due the left atrial contraction. In addition, we measured mitral annular tissue Doppler for both E’ (passive LV filling), and A’ wave (atrial contraction), respectively (**Figure 2B, Table 2**). Calculation of E/E’ and E/A ratios were significantly altered in dKO mice, suggestive of diastolic dysfunction (**Figure 2C, Table 2**). To corroborate the observed cardiac relaxation deficits, we performed micromanometer catheterization on control (wildtype), obscurin and Obsl1 knockouts as well as dKO animals at 9-12 months of age (**Figures 2D, S2A-B; Table 3**). Hearts of dKO mice at baseline displayed no differences in left ventricular systolic contractile function (LV dP/dt_max_), but a significant slowing of relaxation in early diastole (increase in both LV dP/dt_min_ and monoexponential time constant, *Tau*).

**Table 2.** Doppler echocardiography of 5-6 months old male control (CTL) and obscurin/Obsl1 double knockout (dKO) mice. Shown are averages and standard deviation (SD) values. * p < 0.05, ** p < 0.01 vs CTL by Student’s T-test. Abbreviations: MV – mitral valve, TD – tissue doppler, HR – heart rate.

|  | Age<br>(weeks) | MV HR<br>(bpm) | MV E<br>(cm/s) | MV A<br>(cm/s) | MV E/A | MV E'<br>(cm/s) | MV A'<br>(cm/s) | MV E/E' |
| --- | --- | --- | --- | --- | --- | --- | --- | --- |
| CTL<br>(n=6) | 19.5±1.9 | 273±84 | 38.8±16.6 | 22.7±16.2 | 2.3±1.5 | 1.58±0.52 | 0.85±0.54 | 24.3±4.7 |
| dKO (n=9) | 21.5±4.2 | 333±44<br>(p=0.083) | 50.9±11.1<br>(p=0.099) | 46.4±8.8** | 1.1±0.1* | 1.43±0.43 | 1.51±0.32* | 40.6±13.9* |

**Table 3.** Hemodynamics analysis of control, obscurin knockout, Obsl1 knockout and dKO male mice. Shown are averages and standard error of mean (SEM) values. * p < 0.05, ** p < 0.01, *** p < 0.001 by two-way ANOVA analyses, followed by Dunnett’s multiple comparisons test.

|  | Dobutamine<br>(Units) | Heart rate<br>(bpm) | Max dP/dt<br>(mmHg/s) | Min dP/dt<br>(mmHg/s) | Exponential<br>Tau (ms) | Max Pressure<br>(mmHg) | EDP (mmHg) |
| --- | --- | --- | --- | --- | --- | --- | --- |
| CTL<br>(n=9, average<br>age=14.7<br>months) | 0 (Basal) | 328±16 | 9129±661 | -7004±475 | 13.22±0.71 | 115.3±6.4 | 4.31±0.32 |
|  | 0.75 | 342±15 | 9472±581 | -7435±369 | 13.06±0.68 | 121.3±6.2 | 4.64±0.28 |
|  | 2 | 361±13 | 11345±631 | -8530±320 | 12.52±0.66 | 134.8±6.9 | 5.59±0.34 |
|  | 4 | 409±11 | 14402±473 | -10339±373 | 10.40±0.40 | 137.4±4.3 | 7.30±0.73 |
|  | 6 | 478±11 | 16544±441 | -11836±485 | 8.49±0.37 | 132.7±3.6 | 8.95±0.90 |
|  | 8 | 519±12 | 16030±771 | -12125±674 | 7.67±0.31 | 125.9±3.6 | 10.86±0.98 |
| Obsc<br>(n=7, average<br>age=14.7<br>months) | 0 (Basal) | 367±15 | 8075±515 | -6074±546 | 12.92±0.46 | 97.9±6.2 | 4.82±0.88 |
|  | 0.75 | 380±15 | 9067±781 | -6542±665 | 13.10±0.49 | 105.8±8.8 | 5.85±1.23 |
|  | 2 | 400±18 | 10934±984 | -7527±772 | 12.54±0.60 | 118.2±9.9 | 7.15±1.70 |
|  | 4 | 443±24 | 14950±1048 | -9551±937 | 11.45±1.38 | 126.0±7.4 | 8.87±1.44 |
|  | 6 | 496±30 | 17012±1184 | -11343±925 | 8.83±0.66 | 128.1±6.3 | 11.02±1.54 |
|  | 8 | 536±27 | 16977±1238 | -12033±946 | 8.03±0.75 | 123.9±5.9 | 13.43±1.43 |
| Obsl1<br>(n=9, average<br>age=13.8<br>months) | 0 (Basal) | 346±15 | 9204±568 | -6850±515 | 13.28±0.76 | 114.8±8.4 | 3.64±0.37 |
|  | 0.75 | 354±14 | 9809±487 | -7491±405 | 13.03±0.86 | 123.4±8.4 | 3.86±0.35 |
|  | 2 | 372±15 | 11726±610 | -8635±442 | 12.27±0.88 | 138.2±9.4 | 4.53±0.46 |
|  | 4 | 414±18 | 15070±735 | -10634±578 | 10.12±0.68 | 147.7±7.7 | 5.80±0.74 |
|  | 6 | 480±12 | 18152±842 | -13419±417 | 8.14±0.38 | 149.8±6.7 | 7.76±0.63 |
|  | 8 | 525±12 | 18064±1216 | -13843±745* | 7.39±0.31 | 138.9±8.3 | 8.41±0.96 |
| dKO<br>(n=6, average<br>age=13.7<br>months) | 0 (Basal) | 331±18 | 7455±866 | -4042±608** | 20.02±1.43*** | 98.0±9.9 | 6.29±0.64 |
|  | 0.75 | 347±16 | 8164±908 | -4575±677** | 19.40±1.54** | 106.8±10.7 | 7.68±0.72* |
|  | 2 | 371±17 | 9173±1154 | -5140±880*** | 19.68±2.03*** | 114.5±13.2 | 9.09±1.06* |
|  | 4 | 435±15 | 10179±1435** | -5555±1129*** | 22.41±4.06*** | 114.3±14.3 | 10.70±1.62* |
|  | 6 | 476±22 | 10484±1655*** | -5799±1189*** | 21.29±4.32*** | 106.8±12.9* | 11.40±1.72 |
|  | 8 | 514±26 | 10860±1642*** | -6006±1156*** | 17.39±3.41*** | 100.5±10.4* | 10.39±2.30 |

Animals in all groups were able to increase heart rates to a similar extent when challenged with the adrenergic receptor agonist dobutamine. However, at higher doses, dKO mice manifested reduced augmentation of LV dP/dt_max_ in response to increasing doses of dobutamine. Altogether, these data indicate that dKO hearts display both a significant slowing of relaxation as well as a loss of myocardial contractile reserve when pharmacologically challenged.

Analysis of pressure volume (P/V) loops revealed an increase in the alpha coefficient of fitted end-diastolic pressure volume relationship (EDPVR) curves, albeit with a p-value of 0.089 (**Figure 2E, bottom left panel; Table 4**). In addition, *Tau* values as calculated by linear least squares fit of dP/dt vs. P during isovolumic relaxation showed again significant increases in dKO hearts (**Figure 2E, bottom middle panel; Table 4**). We also observed on average a greater resting left ventricular end diastolic pressure (LVEDP) in dKO compared to control mice (**Figure 2E, bottom right panel**), suggestive also of increased venous pressures in the pulmonary circulation, which may lead to the development of heart failure cells (siderophages). Indeed, analysis of lung sections stained with Prussian blue revealed the presence of siderophages in dKO lungs (**Figure 2F**). Taken together, our data indicate that hearts of dKO mice display signs of diastolic heart failure.

**Table 4.** Pressure-volume (P/V) loop analysis of 9 months old male wildtype (CTL) and obscurin/Obsl1 double knockout (dKO) hearts. Shown are averages with standard deviation (SD); n=8 for analysis of bolus infusion. *** p < 0.001 vs CTL by two-way ANOVA analyses, corrected for multiple comparisons using Tukey test; or parametric Welch’s T-test.

| Parameter | Volume (μl) of step bolus infusion | CTL | dKO |
| --- | --- | --- | --- |
| Max dP/dt (mm Hg/s) | 0 | 7022±1989 | 7697±1354 |
|  | 250 | 7349±879 | 7112±931 |
|  | 375 | 7083±823 | 6958±864 |
|  | 500 | 7220±801 | 6723±890 |
|  | 750 | 6748±1119 | 6219±702 |
|  | 1000 | 6851±785 | 6195±930 |
| Min dP/dt (mm Hg/s) | 0 | -6825±2058 | -5676±1147*** |
|  | 250 | -7036±1011 | -5356±999*** |
|  | 375 | -6579±1098 | -5193±921*** |
|  | 500 | -6701±1177 | -4990±899*** |
|  | 750 | -6204±1456 | -4555±649*** |
|  | 1000 | -6222±1279 | -4481±859*** |
| Max Pressure (mm Hg) | 0 | 103.4±23.8 | 103.0±11.4 |
|  | 250 | 108.5±14.2 | 102.3±10.1 |
|  | 375 | 108.6±16.2 | 102.5±9.5 |
|  | 500 | 109.7±14.1 | 101.5±9.3 |
|  | 750 | 108.1±17.0 | 99.0±7.4 |
|  | 1000 | 109.8±14.9 | 99.1±6.4 |
| Linear *Tau* from dP/dt vs P (ms) | 0 | 7.76±0.74 | 11.21±1.40*** |
|  | 250 | 7.89±1.12 | 11.68±1.41*** |
|  | 375 | 8.33±1.10 | 11.76±1.37*** |
|  | 500 | 8.18±1.02 | 11.88±1.44*** |
|  | 750 | 8.56±1.04 | 11.96±2.16*** |
|  | 1000 | 8.61±1.06 | 12.96±1.72*** |
| Max dP/dt slope calculated as linear regression from bolus infusion Max dP/dt | - | -0.317±0.878 | -1.233±0.834<br>(p=0.051) |

### Diastolic dysfunction in dKO mice is not accompanied by fibrosis and upregulation of the fetal gene program

Even though diastolic heart failure is difficult to diagnose, and no reliable markers have been reported, this type of cardiomyopathy is often associated with increased levels of natriuretic peptide B (BNP) and fibrosis ^65–68^. Immunofluorescence, immunoblot and hydroxyproline analyses did not detect any interstitial fibrosis or elevated expression of collagen-I and smooth muscle actin (sm-actin) in dKO hearts when compared to controls (**Figures 3A-D**).

**Figure 3.**
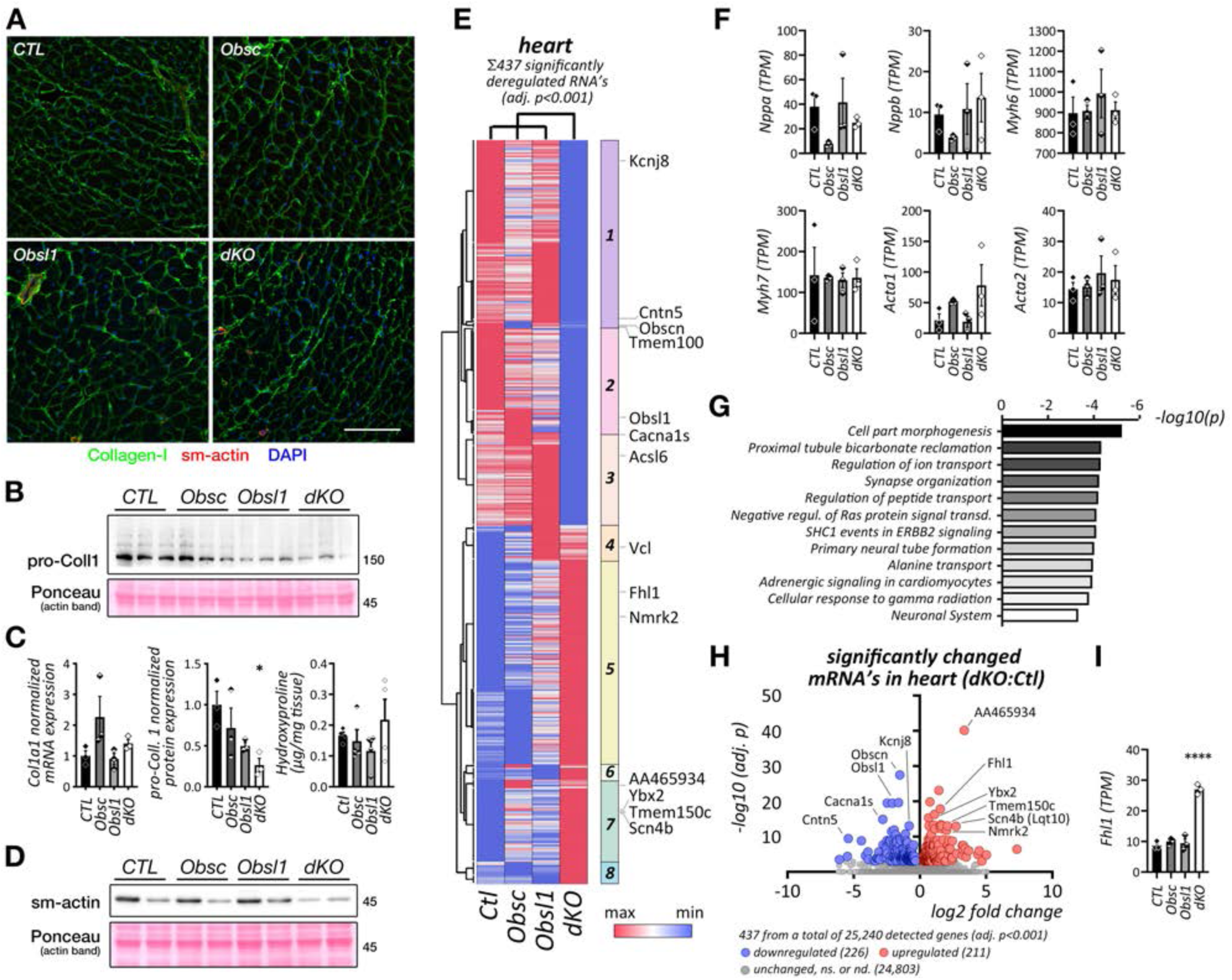
Analysis of cardiac fibrosis and re-expression of the fetal cardiac gene program. A. Immunofluorescence analysis of frozen cardiac sections from wildtype (CTL), obscurin (Obsc) and Obsl1 knockouts as well as obscurin/Obsl1 double knockout animals (dKO), decorated with antibodies against collagen-1 (green) and smooth muscle actin (red). DAPI was used as a counterstain. Scalebar = 100µm. B. Immunoblot analysis of pro-collagen-1 expression levels in all groups of animals. C. Quantification of cardiac collagen-1 mRNA levels (left panel), pro-collagen-1 protein levels (middle panel; normalized to total actin) and total hydroxyproline levels (right panel) in all groups of animals. Shown are averages with standard error of mean values. * p < 0.05 vs. CTL by ANOVA, with multiple comparisons using Dunnett’s test. D. Immunoblot analysis of smooth muscle actin (sm-actin) protein levels in all groups of animals. E. Unbiased analysis of total cardiac mRNA expression by RNA-Seq. Shown is the hierarchical clustering of identified significantly deregulated mRNAs between all groups of animals (adjusted p < 0.001). Representative mRNAs for each of the identified 8 major clusters are highlighted. F. Analysis of cardiac fetal gene program (in transcripts per million, TPM) extracted from the RNA-Seq dataset. Shown are averages with standard error of mean values. G-H. Pathway enrichment analysis and volcano plot of significantly deregulated transcripts from all groups in the RNA-Seq dataset. I. Analysis of Fhl1 expression levels (in transcripts per million, TPM) from the RNA-Seq dataset. Shown are averages with standard error of mean values. **** p < 0.0001 vs. CTL by ANOVA, with multiple comparisons using Dunnett’s test.

To further investigate changes to the gene expression profile we performed RNA-Seq analysis, identifying mRNAs of 437 genes that were significantly differentially regulated between all groups of mice (**Figure 3E**). Our analysis revealed that mRNA expression of fetal gene markers *Nppa* and *Nppb* (encoding for atrial natriuretic peptide (ANP) and BNP, respectively), alpha and beta myosin heavy chain (*Myh6* and *Myh7*, respectively), as well as of pro-fibrotic genes *Acta1* and *Acta2* (encoding for skeletal and smooth muscle actin isoforms, respectively) were unaltered in all sets of mice analyzed (**Figure 3F**). Gene ontology analysis also showed no enrichment for genes involved in fibrotic or inflammatory processes in dKO hearts, observing only significant involvement of genes in ‘regulation of ion transport’ or ‘adrenergic signaling in cardiomyocytes’ that have well-known functions in the heart (**Figure 3G**). Interestingly, our RNA-Seq dataset revealed significant deregulation of several genes associated with cardiac functions (**Figure 3H**), such as upregulation of four and-a-half LIM domain protein *Fhl1* mRNA (**Figure 3I**)^69–71^, deregulation of several genes involved in ion homeostasis and calcium cycling (e.g. *Cacna1s*^72^, *Kcnj8*^73^ or *Scn4b*^74^), or reduction of the mRNA levels for the fatty acid metabolic enzyme *Acsl6*, which was recently linked to diastolic dysfunction^75^.

In summary, these data indicate that the diastolic dysfunction observed in dKO may not result from decreased compliance of the heart due to increased extracellular matrix deposition and fibrosis. In addition, we did not find a change to the fetal-gene program, including increased expression of *Nppb*, which has been reported in some diastolic heart failure studies as a potential biomarker^65, 66^. However, our data suggest that diastolic dysfunction in dKO hearts may be impacted by alterations to cardiac ion transport.

### Obscurin/Obsl1 regulates SR architecture and SR proteins

Obscurin was shown to regulate SR architecture through its interaction with the muscle-specific small ankyrin 1 isoform (sAnk1.5), which is embedded in the SR membrane ^17, 20–23^. No role for Obsl1 has been established for SR architecture and function, as it lacks the sAnk1.5 binding domain found in the C-terminus of obscurin isoform A^22^.

Our analyses showed loss of sAnk1.5 expression and localization at M-bands in obscurin knockout hearts (**Figures 4A-C**), as previously reported for skeletal muscles^23^. Protein levels and localization of sAnk1.5 in hearts of dKO mice were indistinguishable when compared to obscurin knockouts. Surprisingly however, there was a significant reduction in sAnk1.5 protein levels and impaired localization at M-bands in Obsl1 knockout hearts, a finding that is contrary to our observations in skeletal muscle-specific Obsl1 knockouts^43^.

**Figure 4.**
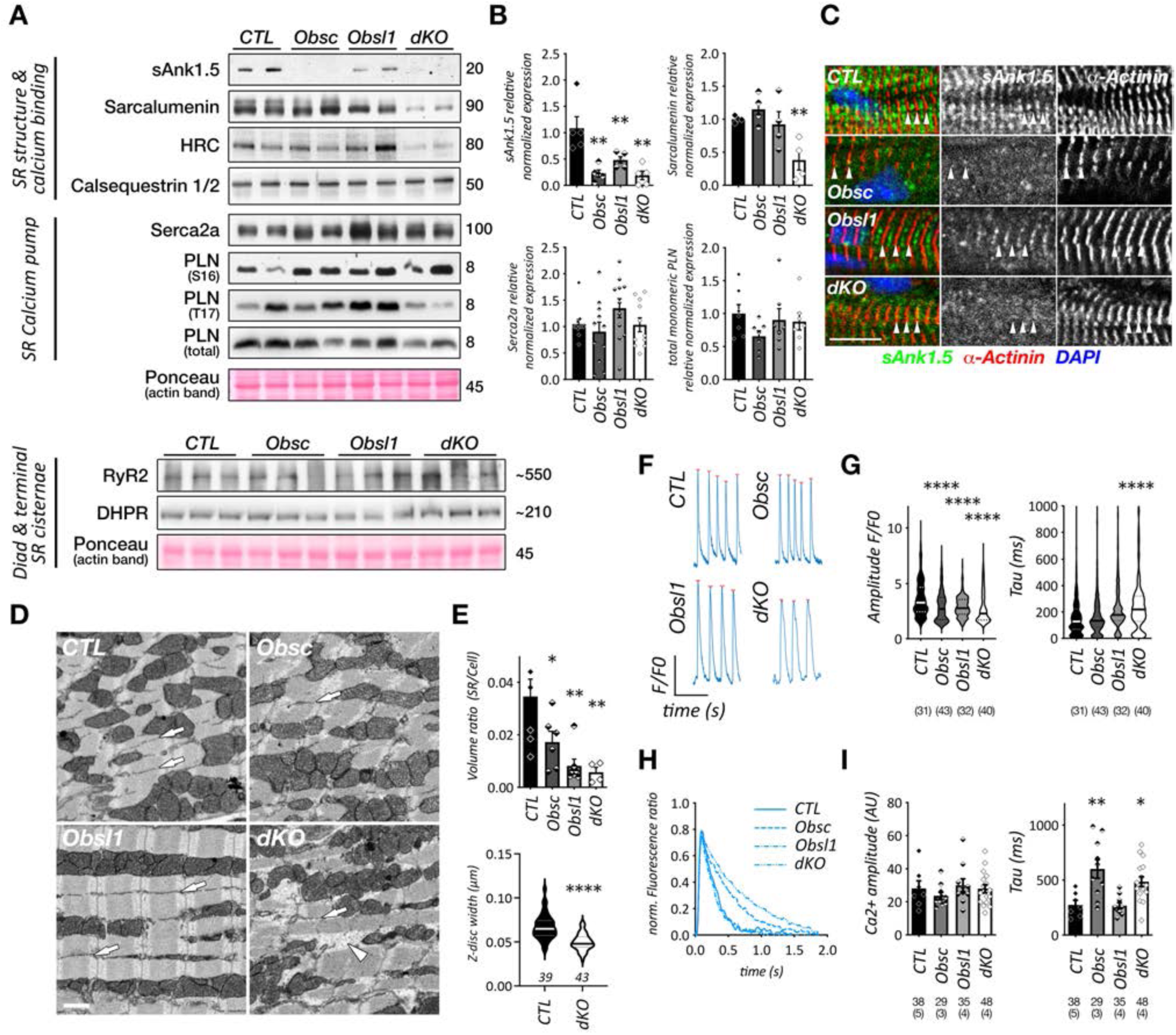
Lack of obscurin/Obsl1 alters sarcoplasmic reticulum architecture and calcium cycling. A. Immunoblot analysis of whole cardiac extracts from wildtype (CTL), obscurin (Obscn) and Obsl1 single knockout, as well as obscurin/Obsl1 double knockout (dKO) animals. Ponceau stained actin is shown as loading control. B. Quantification of relative small ankyrin 1.5 (sAnk1.5), sarcalumenin, Serca2a and phospholamban (PLN) levels normalized to total actin. Shown are averages with standard error of mean values. ** p < 0.01 vs. CTL by ANOVA, with multiple comparisons using Dunnett’s test. C. Immunofluorescence of cardiac sections from all groups stained with antibodies against sAnk1.5 (green in the overlay) and sarcomeric ⍺-actinin 2. DAPI was used as a counterstain. Arrowheads highlight location of sarcomeric Z-discs. Scalebar = 10µm. D. Representative images from serial block face electron micrographs (SBEM) image stacks of cardiac samples from all groups of animals. Arrows highlight sarcoplasmic reticulum, while arrowheads indicate areas of cytoplasm in dKO cardiomyocytes devoid of mitochondria and sarcomeres. Scalebar = 0.5µm. E. Analysis of SR volume to cell volume ratio and Z-disc widths from SBEM image stacks. Shown are averages with standard error of mean values. (E, top panel) or median (solid line) and quartiles (dashed lines) in violin plots (E, bottom panel). Sample sizes are indicated in the figure. * p < 0.05, ** p < 0.01, ** p < 0.001, p < 0.0001 vs CTL. by ANOVA, with multiple comparisons using Dunnett’s test (top panel) or T-test (bottom panel). F-G. Analysis of cardiac calcium cycle in spontaneously beating cultures of neonatal mouse cardiomyocytes from all groups of animals. Representative calcium traces (F) and subsequent analysis for changes to amplitude (G left panel) and Tau (G right panel). Sample sizes of analyzed cells (N) are indicated in the figure. Cells were isolated from at least two litters (4-5 neonates) per group. Shown are median values (solid line) and quartiles (dashed line) in violin plots. **** p < 0.0001 by ANOVA, with multiple comparisons using Dunnett’s test. H-I. Calcium transients of freshly isolated adult cardiomyocytes from 6 months old animals from all groups. Representative images of calcium traces (H) and subsequent quantification of calcium amplitudes and Tau values. Sample sizes of biological replicates (in brackets) and analyzed cells are indicated in the figure. Shown are averages with standard error of mean values. * p < 0.05, ** p < 0.01 vs CTL by ANOVA.

These changes in sAnk1.5 were not mirrored in changes to Serca2a or phospholamban (Pln) expression (**Figures 4A-C**). Protein levels of the ryanodine receptor (RyR) and L-type calcium channels (Dhpr) remained unchanged. We also determined alterations to calcium binding proteins located within the SR lumen. Protein levels of calcium binding proteins sarcalumenin and histidine-rich calcium binding protein (HRC) were significantly decreased in hearts of dKO mice only, indicating that effects of obscurin/Obsl1 loss on SR architecture and function are exacerbated in the dKO mice (**Figures 4A-B**). Intriguingly, levels of calsequestrin, another calcium binding protein located at the junctional SR lumen, remained unaltered between all investigated groups.

### Lack of obscurin/Obsl1 modulates cardiac SR and calcium cycling

To gain insights on how a lack of obscurin and/or Obsl1 changes muscle structure and SR architecture in high resolution, we employed serial block face transmission electron microscopy (SBEM) using cardiac samples from 9-month-old wildtype control, obscurin knockout, Obsl1 knockout and dKO mice (**Figures 4D-E**). Three-dimensional reconstruction of EM-stacks allowed for precise outlining of the SR (**arrows in Figure 4D**), and measurement of SR volumes, which were significantly reduced for all single knockouts and dKO hearts (**Figure 4E, top graph**). While we did not observe any overt differences in sarcomeric structure, analysis of Z-disc widths showed a significant reduction in dKO hearts (**Figure 4E, bottom graph**). We also noticed the occasional appearance of cytoplasmic regions void of sarcomeres that are in close association to mitochondria in dKO cardiomyocytes only (**arrowhead in Figure 4D**).

To test if changes to the SR affect cellular calcium cycling, we isolated neonatal and adult cardiomyocytes from all groups. Analysis of calcium transients from spontaneously beating neonatal cardiomyocytes showed decreased calcium amplitudes for all single knockouts and dKO cells. Only cardiomyocytes from dKO hearts displayed significantly increased calcium reuptake times (*Tau*) (**Figures 4F-G**). This increase in *Tau* values was also observable when analyzing calcium transients in freshly isolated and paced adult dKO and obscurin knockout cardiomyocytes (**Figures 4H-I**). Adult cardiomyocytes however showed no changes to calcium amplitudes.

### Metabolic changes in Obsc/Obsl1 dKO mouse hearts indicate a metabolic switch to glycolysis

To gain unbiased insights into changes to knockout hearts, we performed proteome analysis (**Figures 5A-B**). Enrichment analysis of significantly altered proteins in dKO hearts suggested primarily changes to cellular metabolism, including the glycolysis pathway (**Figures 5C-D**).

**Figure 5.**
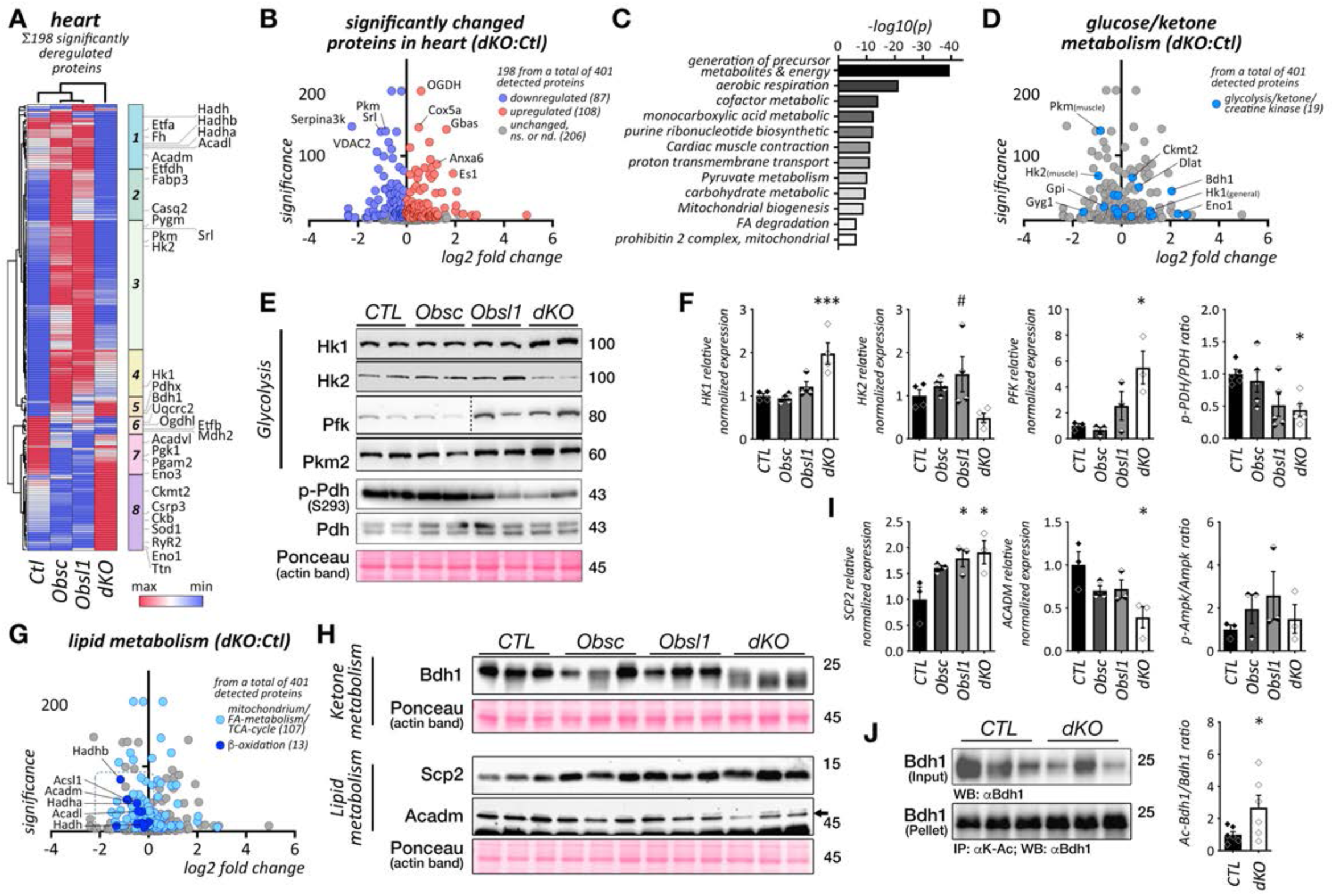
Unbiased proteome analysis shows changes to cardiac metabolism. A-B. Hierarchical cluster analysis and volcano plot of significantly changed protein expression levels from cardiac extracts of wildtype (CTL), obscurin (Obsc) and Obsl1 knockout, as well as obscurin/Obsl1 double knockout (dKO) hearts. Representative proteins for each of the identified 8 major clusters are highlighted (in A). C. Pathway enrichment analysis of significantly deregulated proteins. D. Volcano plot of significantly deregulated metabolic proteins involved in glycolysis, ketone and phosphocreatine metabolism. E-F. Immunoblot analysis I and quantification (F) of glycolysis pathway enzymes hexokinase 1 and 2 (Hk1 and Hk2), phosphofructokinase (Pfk) and pyruvate kinase (Pkm2) as well as total and phosphorylated levels of pyruvate dehydrogenase (Pdh). Ponceau stained actin is shown as loading control. Shown are averages with standard error of mean values, normalized to total actin (in F). * p < 0.05, *** p < 0.01 vs CTL, or # p < 0.05 vs. dKO by ANOVA, with multiple comparisons using Dunnett’s test. G. Volcano plot of significantly deregulated mitochondrial proteins (light blue), specifically highlighting proteins involved in beta-oxidation (dark blue). H-I. Immunoblots (H) and quantification of results (I) of proteins involved in ketone and lipid metabolism. Ponceau stained actin is shown as loading control. Shown are averages with standard error of mean values, normalized to total actin (in I). * p < 0.05 vs CTL by ANOVA, with multiple comparisons using Dunnett’s test. J. Immunoblots (left) and quantification (right) of acetylated Bdh1 immuno-precipitates to total Bdh1 ratios from cardiac extracts of wildtype (CTL) and dKO hearts. Shown are averages with standard error of mean. * p < 0.05 vs CTL by T-test.

Immunoblot assays for rate-determining glycolytic enzymes hexokinase (Hk), phosphofructokinase (Pfk) and pyruvate kinase (Pkm) revealed increased levels of Hk1, Pfk and Pkm2, while Hk2 appeared downregulated in dKO hearts when compared to controls (**Figure 5E-F**). This result, and the significant downregulation of proteins related to beta-oxidation and fatty acid transport observed in our proteome analyses (e.g. Acadl, Acadm, Hadh, Etfa; **Figures 5A, 5G-H**) indicates that dKO hearts may metabolically switch from burning fatty acids to utilizing glucose for energy generation. Indeed, severely downregulated phosphorylation levels of pyruvate dehydrogenase (Pdh) found in dKO hearts (**Figure 5E-F**) strongly suggests increased usage of the glycolysis pathway, rather than utilization of beta-oxidation^76–78^. We also noticed deregulated levels of 3-hydroxybutyrate dehydrogenase 1 (Bdh1), a key enzyme in ketone metabolism in dKO hearts^79^. Immunoblot analyses of Bdh1 indicated a change in the migration pattern of the protein in dKO hearts only, which may be associated with increased lysine-acetylation of Bdh1 (**Figures 5H-J**).

### Mitochondrial size and cristae structure are affected in dKO hearts

In the analyses of our SBEM images we also noted a significant reduction in mitochondrial volume in Obsl1 knockout and dKO mouse heart cells (**Figure 6A**). The reduction in mitochondrial volume indicated a fusion or fission deficit caused by loss of Obsl1. We investigated proteins important for mitochondrial fusion (e.g. mitofusins) or fission (Drp1). While no significant changes were observed for mitochondrial fusion proteins mitofusin-1 (Mfn1) and mitofusin-2 (Mfn2), protein and phosphorylation levels of Drp1 were significantly altered in dKO mouse hearts (**Figure 6B-C**).

**Figure 6.**
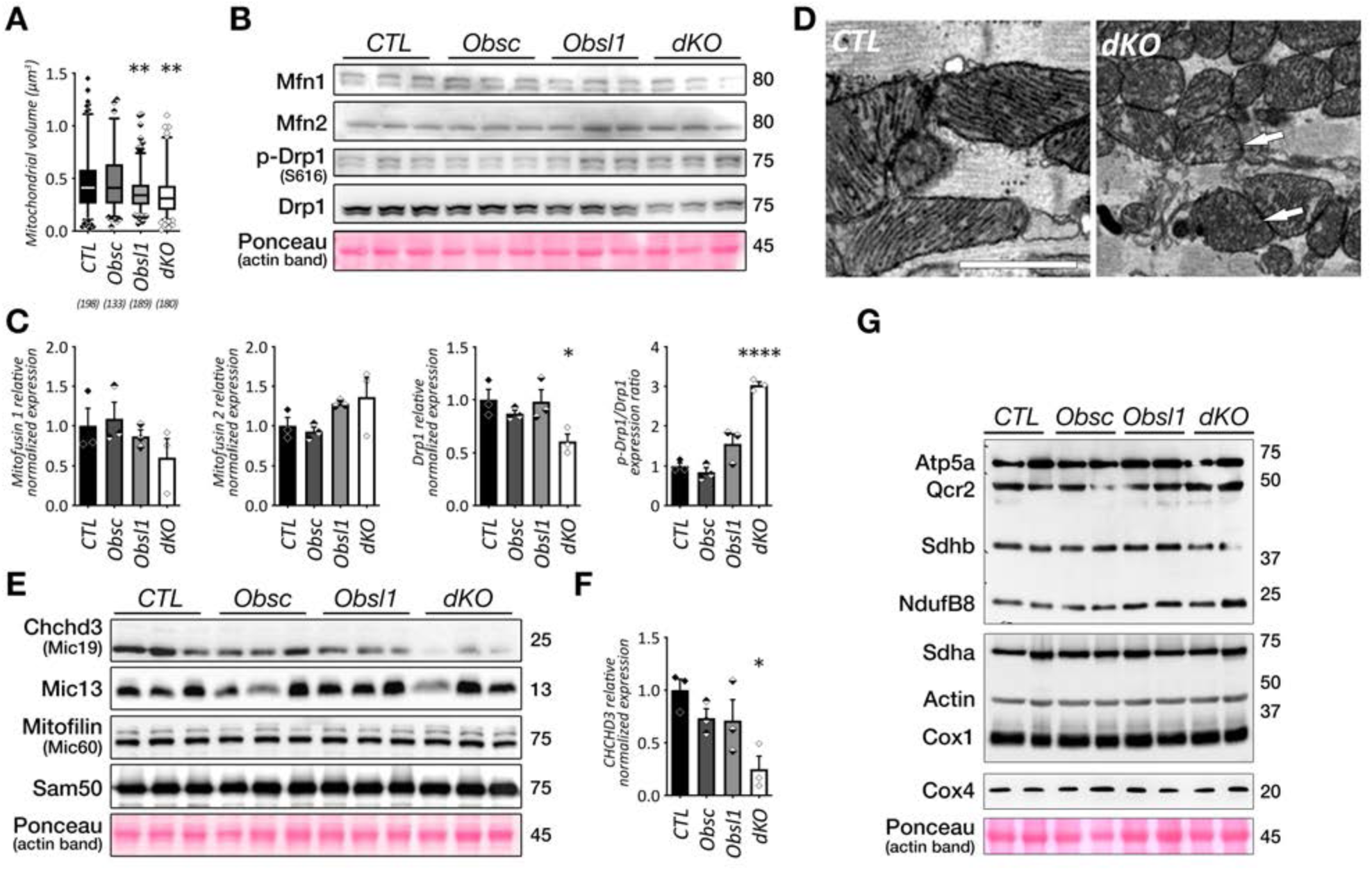
Analysis of cardiac mitochondrial phenotype. A. Quantification of mitochondrial volumes from SBEM image stacks of hearts from wildtype (CTL), obscurin (Obsc) or Obsl1 knockouts as well as obscurin/Obsl1 double knockout (dKO) animals (see Figure 4D). Shown are box and whisker plots with averages (solid line) and 5-95 percentiles. Sample sizes are indicated in the figure. ** p < 0.01 vs CTL by ANOVA, with multiple comparisons using Dunnett’s test. B-C. Immunoblot analysis (B) and quantification (C) of mitofusin (Mfn) protein levels as well as total and phosphorylated dynamin related protein 1 (Drp1) levels in cardiac extracts from all groups. Ponceau stained actin is shown as loading control. Shown are averages with standard error of mean normalized against total actin or ratio of phosphorylated to total protein. * p < 0.05, **** p < 0.0001 vs CTL by ANOVA, with multiple comparisons using Dunnett’s test. D. Representative electron micrographs of mitochondria in hearts of CTL or dKO mice. Arrows indicate mitochondria with disrupted cristae structure. Scalebar = 200nm. E-F. Immunoblot analysis I and quantification (F) of mitochondrial proteins of the MICOS complex involved in cristae formation as well as Sam50 channel. Ponceau stained actin serves as a loading control. Shown are averages with standard error of mean normalized against total actin. * p < 0.05 vs CTL by ANOVA, with multiple comparisons using Dunnett’s test. G. Immunoblot analysis of electron transport chain complex subunits. Ponceau stained actin is shown as loading control.

Our high-resolution analyses of mitochondria also revealed significant disruption of organized cristae structure (**Figure 6D**), suggesting changes to Micos complex proteins^80^. Analysis of select complex components revealed significant loss of Chchd3/Minos19, while other Micos complex proteins were unaffected (**Figures 6E-F**). Chchd3 was recently shown to interact with Obsl1 in a high-throughput binding study^56^ and our knockout data support the mechanistic link to Obsl1. However, only combined loss of Obsl1 and obscurin resulted in a change of Chchd3 protein levels, also suggesting roles for obscurin in regulating protein levels of this Micos complex subunit.

Nonetheless, despite all these changes, analysis of mitochondrial matrix proteins and respiratory chain complexes revealed no significant differences between hearts from all investigated groups (**Figure 6G**), a finding that contrasts with results obtained from skeletal muscle-specific dKO mice^43^.

### Loss of obscurin and Obsl1 alters cellular autophagy and mitochondrial turnover

Our SBEM data of dKO mouse hearts also revealed the occasional appearance of double membraned autophagosomes in close proximity to mitochondria (arrow in **Supplemental Figure S3A**). We wondered, what could be a mechanistic link between obscurin proteins and autophagy. Screening for novel binding partners of the Obsl1 by yeast two-hybrid^43^ identified Atg4d (**Supplemental Figures S3B-C**). Atg4d is a mitochondrially linked^81^ member of the Atg4 cysteine peptidase family involved in cleaving C-terminal ends of LC3, Gabarap or Gabarap-L1 polypeptide precursors, and catalyzing the delipidation of these carrier proteins, e.g. transform lipid carrying Lc3-II to lipid-free Lc3-I^82, 83^.

Biochemical binding assays verified the association of C-terminal Obsl1 Ig-domains 19-20 with Atg4d, also indicating that Obsl1 binding may extend to other members of the Atg4 peptidase family (**Figure 7A**). Presence of Obsl1 Ig19-20 did not affect processivity of Atg4 to cleave pre-Lc3a or pre-Gabarap-L1 (**Supplemental Figures S3D-E**), suggesting that Obsl1 may help localize or direct Atg4 peptidases, rather than modifying their activity.

**Figure 7.**
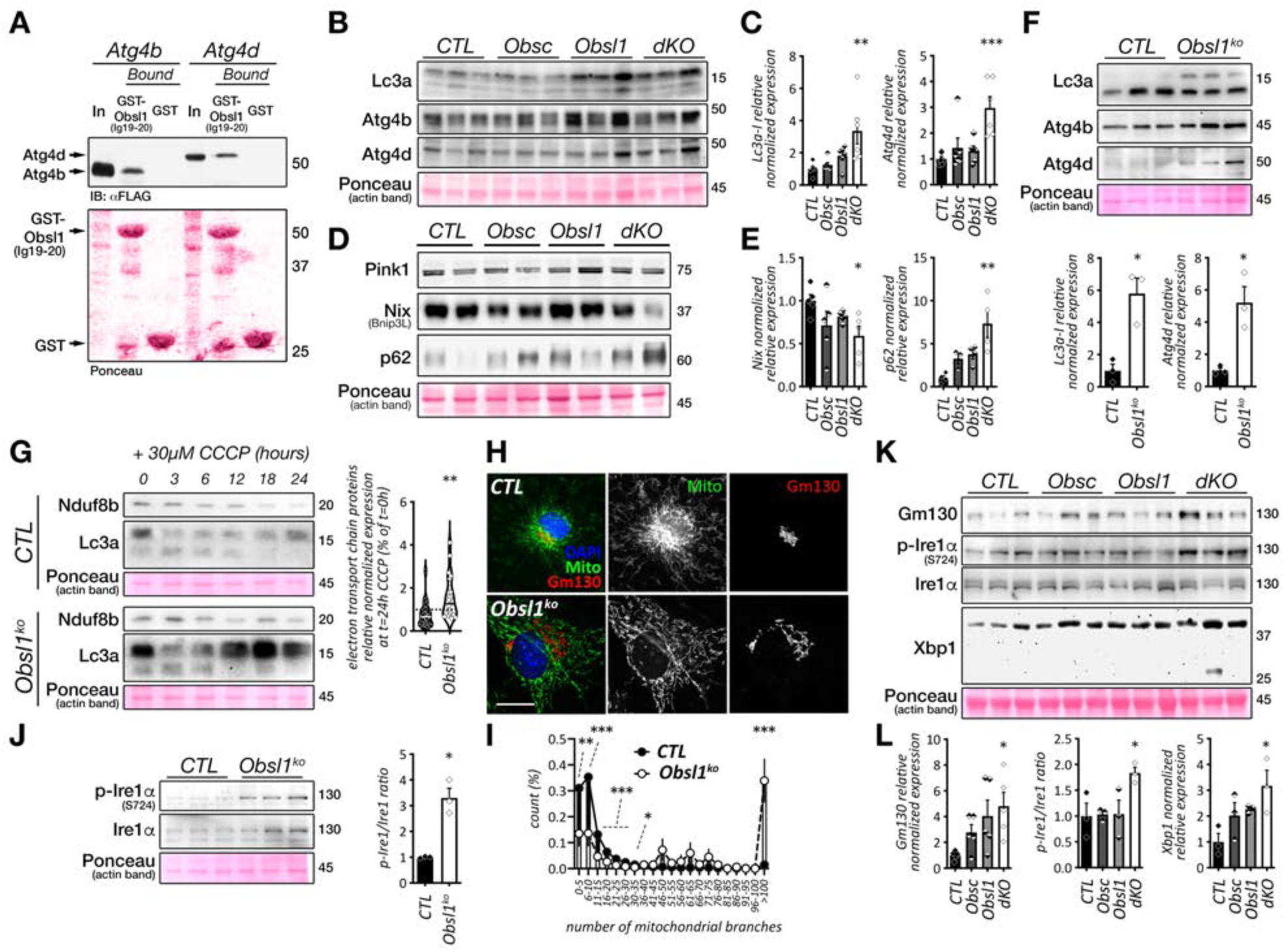
Lack of obscurin and Obsl1 deregulates mitochondrial turnover and results in unfolded protein response stress. A. GST-pulldown assay using either GST or GST fused to C-terminal Obsl1 Ig-domains 19-20 as well as Flag tagged Atg4b and Atg4d. B-C. Immunoblot analysis (B) and quantification (C) of autophagy related Lc3a as well as Atg4b and Atg4d protein levels in cardiac extracts of wildtype (CTL), obscurin or Obsl1 knockout and obscurin/Obsl1 double knockout (dKO) mice. Ponceau stained actin serves as a loading control. Shown are averages with standard error of mean normalized against total actin. ** p < 0.01, *** p < 0.001 vs CTL by ANOVA. D-E. Immunoblot analysis (D) and quantification (E) of Pink1, Nix (Bnip3L) and p62 protein levels in cardiac extracts of wildtype (CTL), obscurin or Obsl1 knockout and obscurin/Obsl1 double knockout (dKO) mice. Ponceau stained actin serves as a loading control. Shown are averages with standard error of mean normalized against total actin. * p < 0.05, ** p < 0.01 vs CTL by ANOVA, with multiple comparisons using Dunnett’s test. F. Immunoblot analysis (top panel) and quantification (bottom panel) of Lc3a as well as Atg4b and Atg4d protein levels in Obsl1 knockout and control (CTL) mouse lung endothelial cells. Ponceau stained actin serves as a loading control. Shown are averages with standard error of mean normalized against total actin. * p < 0.05 by T-test. G. Changes to Nduf8b and Lc3a levels by 30µM CCCP in Obsl1 knockout and control lung endothelial cells analyzed by immunoblots (left panel). Quantification of electron transport chain protein levels (normalized to actin) at 24h CCCP treatment compared to t=0h (dashed line; right panel). Shown are violin plots with median (solid lines) and quartiles. ** p < 0.01 vs CTL by T-test. H-I. Analysis of changes to mitochondria in Obsl1 knockout and control endothelial lung cells by immunofluorescence. Representative images of cells stained with mitotracker (green), Gm130 (red) and DAPI (H). Scalebar = 5µm. Histogram of mitochondrial network branching as determined by skeleton analysis (I). At least 14 cells were analyzed per group. * p < 0.05, ** p < 0.01, ** p < 0.001 vs CTL by multiple unpaired T-test comparisons using false discovery rate (two-stage set-up method of Benjamini, Krieger and Yekutieli). J. Immunoblot analysis (left panel) and quantification (right panel) of total and phosphorylated Ire1⍺ in Obsl1 knockout and control mouse endothelial lung cells. Ponceau stained actin serves as a loading control. Shown are averages with standard error of mean phosphorylated to total protein ratios. * p < 0.05 by T-test. K-L. Immunoblot analysis (K) and quantification (L) of unfolded protein response related Xbp1 protein levels, total and phosphorylated Ire1⍺, as well as Gm130 in cardiac extracts of wildtype (CTL), obscurin or Obsl1 knockout and obscurin/Obsl1 double knockout (dKO) mice. Ponceau stained actin serves as a loading control. Shown are averages with standard error of mean normalized against total actin. * p < 0.05 vs CTL by ANOVA, with multiple comparisons using Dunnett’s test.

We probed Atg4 peptidase protein levels and lipidation of Lc3a and found increased levels of Atg4d, but not Atg4b, as well as elevated levels of de-lipidated Lc3a-I in protein samples of dKO hearts only (**Figures 7B-C**). As Atg4d is specifically linked to mitochondria^81^, we wondered if mitophagy may be affected. Levels of mitophagy-linked Nix/Bnip3L were decreased alongside increased overall p62 levels, suggestive of altered mitophagy specifically in dKO hearts, while Pink and Parkin remained unchanged (**Figures 7D-E, Supplemental Figure S3F**). To further probe the effect of Obsl1 on mitophagy, we tested an established line of murine Obsl1 knockout endothelial lung cells^43, 84^. These cells display similar changes to Atg4d and Lc3-I levels compared to dKO hearts (**Figure 7F**). Induction of mitophagy by depolarizing mitochondria using 30µM CCCP^85, 86^ resulted in decreased degradation of electron transport chain complex proteins, including Nduf8b, Atp5a or Uqcrc2 in Obsl1 knockout cells compared to controls (**Figure 7G, Supplemental Figure S3G**). When analyzing mitochondria in Obsl1 knockout cells by immunofluorescence, we also noticed altered mitochondria structure, with knockout cells displaying a more extensively branched mitochondrial network compared to control cells as determined by skeleton analyses (**Figures 7H-I**). Mitophagy was shown to be interlinked to endoplasmic reticulum (ER) function and deficits in both can induce unfolded protein response (UPR) in cells^87–89^. When analyzing Obsl1 knockout cells, we noticed significantly increased endonuclease inositol-requiring protein 1α (Ire1α) phosphorylation (**Figure 7J**). This result is mirrored in hearts of dKO mice, which also showed increased Ire1α phosphorylation in addition to elevated Xbp1 and Gm130 levels (**Figures 7K-L**).

## Discussion

Members of the obscurin protein family have been demonstrated to play important roles in the function of skeletal muscles. One of the key findings for obscurin was its function for sarcolemma integrity and organization of the SR^20, 24, 25, 43, 90–92^. Less is known for cardiac roles of obscurin and specifically Obsl1. Here, we found that cardiac specific loss of obscurin and Obsl1 results in diastolic dysfunction and heart failure, characterized by shortened lifespans, mitochondrial defects and altered metabolism.

We employed several techniques to characterize the cardiac relaxation problem in dKO mice. Invasive hemodynamics and echocardiography assisted P/V-loop measurements were in our hands the most robust method to measure diastolic functions, as also reported in the literature^93^. Data derived by pulsed wave and tissue Doppler echocardiography, while non-invasive and more practical, may not necessarily by themselves indicate myocardial abnormalities^94^ and require skilled experimenters, as they can be easily impacted by respiratory variation or sample volumes moving in and out of the plane of the annulus. The combined approach, however, clearly indicated impaired cardiac relaxation pattern with mildly elevated filling pressures, possibly accompanied by a slight alteration in left heart compliance as the cause(s) for diastolic dysfunction in dKO mice.

On the molecular level, the cardiac relaxation problem and diastolic dysfunction is brought on by cardiac-specific loss of both proteins, obscurin and Obsl1. Deletion of obscurin alone causes sarcoplasmic reticulum impairment and increased reuptake times for intracellular calcium in diastole. Notable exceptions are the lumenal SR calcium binding proteins sarcalumenin and histidine-rich calcium binding protein, as these are only decreased in hearts of dKO mice. Hence, most changes to the SR and calcium cycling that are also visible in obscurin knockouts may by themselves not be sufficient to result in diastolic dysfunction exhibited by dKO hearts. Loss of Obsl1 alone on the other hand causes mitochondrial fragmentation in cardiomyocytes. However, only the combined loss of both proteins in the heart impacts mitochondrial fission, turnover, unfolded protein response and metabolism. Intriguingly, changes to mitochondrial turnover in dKO hearts were not linked to changes Pink1 or Parkin expression levels. Intriguingly, modulation of mitophagy has been demonstrated to protect against cardiomyopathy, with inhibition of mitophagy leading to exacerbated diastolic and systolic dysfunction^95^. Altered cardiac metabolism and loss of mitochondria cristae structure and function was also specifically linked to diastolic dysfunction development^96^. Mitochondria of diastolic heart failure patients and in a mouse model for heart failure with preserved ejection fraction show cristae destruction and reduced mitochondrial area^97, 98^.

There are several differences between the roles of obscurin and Obsl1 in skeletal muscle and the heart. One of the principal binding partners for the C-terminus of obscurin-A is the muscle specific sAnk1.5 isoform of Ankyrin 1^20–23, 90^. While loss of obscurin alone in skeletal muscles decreased levels of sAnk1.5, deletion of Obsl1 had no effect on sAnk1.5 levels^43^. In heart however, loss of Obsl1 negatively affected sAnk1.5 protein levels and localization (**Figures 4A-C**). This loss of sAnk1.5 in absence of Obsl1 but presence of obscurin suggests an unknown role for Obsl1 either in localizing or stabilizing this small muscle-specific ankyrin isoform in the heart. Another difference is that while proteome analyses show marked changes in electron transport chain proteins, immunoblot analysis using OxPhos cocktail antibodies failed to elicit the downregulation observed in skeletal muscle-specific dKO mice^43^. Indeed, there is only limited overlap when comparing significantly altered proteins identified between hearts and skeletal muscles (tibialis anterior and soleus) from dKO mice (**Supplemental Figure S4; Supplemental File 4**). Only 14 proteins were found in all three dKO tissues, albeit not necessarily with the same directionality of change (ie. down- vs. up-regulated).

There are several other animal models available to study obscurin functions. Obscurin Ig58-59 deletion mice show altered calcium handling and arrhythmias^33^. We also noted in our experiments using isolated adult cardiomyocytes from 6 months old obscurin knockouts and dKO hearts significantly prolonged calcium transients, reflected in increased *tau*-values. However, adult cardiomyocytes of obscurin or Obsl1 knockouts, as well as dKO mice displayed no changes to calcium amplitude, perhaps reflecting a gain of function in mice lacking obscurin Ig58-59. Mice with a deletion in Ig58-59 also showed reduced Pln levels. Recently, a binding site for Pln in obscurin Ig58 was characterized that displayed increased binding capacity when mutated to R4344Q, a potentially pathogenic gene variant identified in a hypertrophic cardiomyopathy patient^32^. However, Pln binding to obscurin is contested^34^. Our data on unchanged Pln levels in the single and double knockout animal models suggest no mechanistic link to this micropeptide important for Serca function. However, altered Pln phosphorylation specifically at Thr17 suggests that further experimental testing may be required. The R4344Q mutant of obscurin in mice also led to increased expression of most metabolic proteins, including enzymes involved in lipid catabolism^99^. Our data indicate that loss of obscurin alone had no pronounced effect on genes involved in lipid metabolism, suggesting that other mechanisms are at play in R4344Q mice, resulting in a deregulated cardiac metabolism. This manuscript also identified increased expression of protein kinase G (Pkg), another protein that has been mechanistically linked to diastolic heart failure development^99^. However, unchanged E/A ratios even in older animals suggest no diastolic dysfunction in R4344Q mice. Intriguingly, there are reports of patients with a bi-allelic loss of obscurin that predispose these individuals to recurrent rhabdomyolysis^35^. Data from our obscurin knockout mice support the finding for the benign cardiac phenotype in affected patients. However, our analyses of altered SR architecture that also impacts cardiac calcium cycling suggests some molecular and physiological changes in hearts of these individuals that may be pathological in certain instances, e.g. also predisposing these individuals to cardiomyopathy in case of additional cardiac insults. Indeed, abnormal intracellular calcium handling has also been identified as a major cause of systolic and diastolic dysfunction^100^.

Loss of Obsl1 on the other hand is associated with the development of 3M syndrome, which is characterized by severe pre- and postnatal growth impairment^57–59, 101^. Global Obsl1 knockouts are embryonic lethal, and have therefore limited use to investigate the syndrome^43^. However, data from knockouts mice and cells suggest that loss of Obsl1 exerts major changes to cellular metabolism, specifically acting on mitochondria. There are no reports on altered mitochondrial structure, function, or turnover in cells of 3M patients. However, a recent manuscript identified a rare genetic variant in *Rmrp*, a long non-coding RNA (lncRNA) and RNA component of the mitochondrial RNA processing endoribonuclease, which causes cartilage hair hypoplasia syndrome (CHH)^102^. This syndrome resembles in some aspects 3M growth syndrome, including short stature of affected individuals. An aspect of *Rmrp* functions is the scavenging of micro RNA’s (miRNAs), and *Rmrp* was shown to have links to cullin mediated turnover, including cullin-7, by regulating Cand1 via miR-766-5p^103^.

To our knowledge, the link between Obsl1 and mitochondrial maintenance has not been previously reported. Binding of Obsl1 to Atg4 proteins Atg4b and Atg4d was mapped to the endopeptidase domain. However, enzymatic function of Atg4b was not affected by Obsl1 in our hands. This suggests either that Obsl1 acts as a linker protein that helps direct substrates to the endopeptidase, or that Obsl1 functions in compartmentalizing Atg4 enzymes. We did not test for an effect of Obsl1 on de-lipidating functions of Atg4 enzymes, which is needed to convert the lapidated forms of Lc3, Gabarap or GabarapL1 into their de-lipidated counterparts. Analysis of Lc3 in dKO hearts and Obsl1 knockout cells only suggests the abnormal accumulation of Lc3a-I, the delipidated form of the protein, but no change to levels of the lipidated Lc3a-II. In addition, while loss of Obsl1 alone in hearts increased Lc3a-I levels, the change became only significant in hearts of dKO mice. Moreover, Atg4d levels only increased significantly in dKO hearts. These findings indicate redundancy in functions between Obsl1 and obscurin, with obscurin taking over some of the functionality in Obsl1 knockout hearts. However, the binding site for Atg4 proteins in obscurin remains to be determined.

Besides the newly identified link of Obsl1 to regulating mitophagy via Atg4d, Obsl1 has been reported to bind to Chchd3 (also known as Mic19)^56^, a subunit of the Micos complex important for cristae formation in mitochondria^80^. Loss of Chchd3 in cells results in abnormal mitochondrial morphology, with mitochondria looking more fragmented and displaying reduced cristae content in ultrastructural analyses^104^. The fragmentation of mitochondria in Chchd3 knockdown cells was accompanied by increased Drp1 levels, suggesting deregulated mitochondrial fission. These findings align well with changes observed in dKO hearts, which also show impaired cristae architecture and altered mitochondrial fission, albeit with a significantly increased ratio of phosphorylated to total Drp1 levels. Loss of Obsl1 alone did not result in significant changes to Chchd3 or Drp1. Only loss of both, obscurin and Obsl1 brought on the observed effects, again suggesting some level of functional redundancy between both members of the obscurin protein family.

Further studies are required to understand the ultimate cause for the diastolic dysfunction in dKO mice. These experiments may shed also light into additional pathophysiological mechanisms that drive diastolic dysfunction and heart failure, such as seen in patients suffering from heart-failure with preserved ejection fraction.

## Materials and Methods

### Animals

Obscurin^23^ and Obsl1 gene targeted mice^43^ were crossed with transgenic mice expressing Cre-recombinase under control of a transgenic Nkx2.5 promoter^60^, allowing for cardiac-specific excision of genes. Controls were Nkx2.5-Cre positive wildtypes. If not explicitly stated, all experiments were done using animals from both sexes, using a Black Swiss background. All biochemical and proteome analyses were performed on animals 6-9 months of age. All procedures with genetically modified animals were approved by the Animal Use and Care Committee (IACUC) of the University of California, San Diego.

### Cell culture

Culture of Cos-1 cells was done using growth medium (10% fetal calf serum, 1% penicillin/streptomycin in DMEM) as previously described^105^. Transfection of cells was performed one day after passaging with a cell confluency of 70% and Lipofectamine 2000 (Invitrogen) as per manufacturer’s instructions.

Isolation and culture of primary mouse lung endothelial Obsl1 knockout (flox/-CMV-Cre+) and control cells (flox/+ CMV-Cre+) was previously described^43^. For visualization of mitochondria, cells were incubated in 200 nM Mitotracker Red CMXRos (Molecular Probes) in DMEM at 37°C for 30 minutes. Following incubation, cells were washed with 1x PBS and fixed in 4% PFA diluted in 1x PBS for 5 minutes at room temperature. Cells were stored in 1x PBS at 4°C until further processing. For analysis of mitophagy, cells were incubated with growth medium containing 30 µM CCCP (carbonyl cyanide m-chlorophenyl hydrazone) the day after plating. Following incubation in presence of CCCP, cells were harvested in a modified version of SDS sample buffer^106^ and run on SDS-page following immunoblot analyses.

### GST Pulldown

To test binding of Atg4 proteins to the C-terminal Obsl1 Ig-domains 19-20, GST pulldown assays were employed ^105^. Briefly, Flag-tagged Atg4 proteins were expressed in transfected Cos-1 cells. Two days following transfection, cells were lysed in IP buffer (150 mM NaCl, 10mM Tris-HCl pH 8, 1 mM DTT, 0.2% NP-40, 1x protease inhibitor [Roche], 1x phosphatase inhibitor [Roche]), sonicated and centrifugated for 10 minutes at 14,000 RPM at 4°C. Following centrifugation, soluble proteins were precleared with Glutathione Sepharose beads (bioPlus fine research chemicals) for 2 hours at 4°C on a rotary shaker. Precleared protein lysates were incubated either with GST-Obsl1-Ig19-20 or GST alone for 2 hours at 4°C on a rotary shaker, before washing of beads for three times using ice-cold wash buffer (1x PBS with 0.2% NP-40). Input and pellet fractions were analyzed on SDS-page following immunoblotting to determine binding of proteins.

### Cloning of bacterial and eucaryotic expression constructs

Obsl1 fragments were subcloned in-frame into a custom GST-C1 vector from GFP-Obsl1 constructs^43^. Eucaryotic expression constructs for human Atg4b (NM_013325), murine Atg4d (NM_153583), human Lc3a (NM_032514) and human GabarapL1 (AF087847) were done by amplifying coding sequences via polymerase chain reaction, followed by in-frame cloning into eucaryotic expression vectors pEGFP-C1 (Clontech) or a custom-built Flag-C1 vector, where the coding sequence of pEGFP was replaced in-frame by the Flag epitope. All constructs were sequenced to determine correct integration into the vector.

### Immunofluorescence and histology

For immunofluorescence, cardiac samples were frozen in OCT medium (Tissue Tek) before being processed by cryosectioning on a Leica cryostat. Frozen sections were dried at room temperature and stored at −80°C until further use. Staining of sections was done as previously described^107^. In short, sections were fixed in acetone at −20°C for 5 minutes, before rehydration at room temperature using 1x PBS for 5 minutes. Following permeabilization with 0.2% Triton X-100 in 1x PBS for 5 minutes at room temperature, sections were blocked with 5% normal donkey serum (Jackson Immunoresearch) and 1% BSA (Sigma-Aldrich) in Gold Buffer (55 mM NaCl, 2 mM EGTA, 2 mM MgCl2, 20 mM Tris-HCl, pH 7.5) for 1 hour in a humid chamber, before incubation with a mix of primary antibodies dissolved in Gold Buffer supplemented with 1% BSA (Sigma-Aldrich) over night at 4°C. Subsequently, sections were washed three times 5 minutes at room temperature with 1xPBS, before incubation with secondary antibodies in Gold Buffer supplemented with 1% BSA for 1 hour in a humid chamber. Sections were washed three times in 1x PBS before mounting in Fluorescent Mounting Medium (DAKO).

Staining of PFA fixed cells with primary and secondary antibodies was processed similarly. Imaging was done using a Fluoview 1000 confocal laser scanning microscope (Olympus), equipped with 10x air or 40x oil objective in sequential scanning mode and Zoom rates between 1 and 4. Images were processed using ImageJ (NIH, version 1.53) with the Bio-Formats plugin^108^ and figures were assembled using Adobe Photoshop (Adobe). Analysis of the mitochondrial network was done using the skeleton analysis tool in ImageJ, as described elsewhere^109^.

For histology, whole hearts or isolated lung lobes were fixed overnight in 4% PFA in 1xPBS at 4°C. Following fixation, hearts were washed in 1x PBS for 1 hour at room temperature, before incubation with 50% OCT in 1x PBS solution overnight at 4°C. Subsequently, hearts were frozen in OCT and sectioned on a cryostat (Leica). Sections were stained with hematoxylin and eosin stain kit (Sigma Aldrich) as per manufacturer’s instructions and imaged on an Olympus SZX12 stereomicroscope equipped with a Lumenera Infinity 3 CCD camera and the Infinity Capture Software (Version 6.4.0; Lumenera).

Lungs were embedded in paraffin and sectioned as previously described^110^. For staining of heart failure cells, sections were deparaffinized, and incubated for 20 minutes in a freshly made solution containing 10% hydrochloric acid and 5% Potassium ferrocyanide, Trihydrate (Sigma Aldrich) in distilled water. Following staining, sections were washed for three times with distilled water before dehydration using increasing amounts of ethanol, two washes with Histoclear (VWR) and embedding in Permount (Thermo Fisher). Stained sections were imaged in brightfield mode on a Revolution Microscope (Echo), equipped with a 10x and 20x air objective (Olympus).

### Isolation of neonatal mouse cardiomyocytes and analysis of calcium transients

Isolation and culture of neonatal mouse cardiomyocytes was done as previously described^111^. Briefly, newborn mice were euthanized and hearts were dissociated by enzymatic digest. Genotyping was done to confirm the desired genotype for all subsequent experiments. Digested hearts were pooled and cells were preplated onto 10cm dishes with plating medium^111^. Following 2 hours of incubation, non-adherent cardiomyocytes were washed off the plate and finally plated into 35mm glass bottom dishes (Ibidi). Cells were switched to maintenance medium the following day and cultured for two more days.

For imaging of calcium transients, cells were moved into HBSS medium (Thermo Fisher), supplemented with glucose to a final concentration of 25mM. Cells were loaded with Fluo-4 AM (Thermo Fisher) for 20 minutes, washed with HBSS supplemented with glucose and imaged using line scans on an inverted confocal laser scanning microscope, equipped with a 40x oil objective and zoom rates between 1x and 4x(Olympus Fluoview 1000). Traces of spontaneously beating neonatal cardiomyocytes were subsequently quantified using ImageJ (NIH, version 1.53) and analyzed using a custom script (**Supplemental File 2**) in Matlab (version R2021a).

### Isolation of adult ventricular myocytes

Single ventricular myocytes were obtained from adult mouse hearts by enzymatic dissociation^112^. Briefly, hearts were excised and perfused through the coronary arteries on a Langendorff system with a calcium-free physiological saline solution followed by a Ca^2+^/enzyme-containing saline mixture (Type II collagenase and protease XIV; Worthington and Sigma Aldrich, respectively). Single isolated cells were separated by mechanical agitation and cellular debris was removed by filtration through a nylon mesh with 250-μm porosity. Fine cellular debris was minimized as healthy myocytes were allowed to sediment by gravity, and supernatant solution was removed, at the end of successive stepwise increases of calcium concentration to reach 1.8 mM. Cells were plated on laminin coated dish using plating medium^112^ and observed within 16-20 hours following cell culture.

### Contractility and calcium measurements in adult cardiac myocytes

Sarcomere shortening and intracellular calcium transients were measured using an IonOptix recording system 16h after the myocytes were isolated. Calcium transients were measured from myocytes paced at 1Hz and loaded with 2 μM Fura2-AM (Molecular Probes) for at least 20 min at 37°C (allowed de-esterification period) following a 20 min wash step with DMEM. After the wash step, coverslips were placed on an inverted microscope and perfused with Tyrode’s solution (137 mM NaCl, 0.5 mM MgCl2, 5 mM Glucose, 4 mM KCl, 1.8 mM CaCl2, 10 mM HEPES, pH 7.4) at room temperature. All cover slips were used within 90 min of loading. Changes in sarcomere shortening were recorded in real time with a high-speed CCD camera (Ionoptix). Cellular fluorescence emission wavelengths (510 nm) were recorded using a photomultiplier tube from myocytes excited with 340 nm (calcium independent measure) and 380 nm (calcium dependent measure) excitation wavelengths. Only myocytes with a resting sarcomere length greater than 1.9 μm that followed the field stimulation protocol were included in the study. Ionwizard software (IonOptix LLC, Milton, MA) was used to analyze the results and quantify the data. Velocity, magnitude, relaxation times, sarcomere shortening and calcium release were used to quantify the features of each group of cells.

### RNA expression analysis and RNA-Seq

RNA isolation from whole heart extracts was done as previously described^107^. In short, total cardiac RNA from three hearts per group was isolated using Trizol (Ambion) as per manufacturer’s instructions. Resulting RNA was snap frozen and stored at −80°C until qRT-PCR analysis or library generation for RNA-Seq analyses. Expression levels of *Collagen1a1* mRNA from total cardiac RNA was determined using PerfeCTa SYBR green real-time qPCR mix (Quanta BioSciences) and a CFX96 thermocycler (Bio Rad) as described elsewhere^113^, using the following set of oligonucleotides^114^: Col1a1.fwd TCACCAAACTCAGAAGATGTAGGA and Col1a1.rev GACCAGGAGGACCAGGAAG. For RNA-Seq analysis, the sequencing library was prepared using a TruSeq stranded mRNA library preparation kit with polyA selection (Illumina). 150 cycles paired-end sequencing were performed using the NovaSeq 6000 system and v1 sequencing chemistry (Illumina) aiming for average coverage of 30 M reads per sample. Next generation RNA-Seq was done by the SNP&SEQ Technology Platform (Uppsala, Sweden). Bioinformatics processing was done as described elsewhere^110^. Briefly, quality of the reads was assessed using MultiQC^115^, sequences were aligned using STAR^116^ and read counts were summarized using featureCounts^117^. The DESeq2 package ^118^ running in R (v. 4.1.0, https://www.R-project.org/) was used to assess differentially expressed genes. FDR correction was applied to adjust p values. Comparisons with log2 fold change >0.3 or <-0.3, and p < 0.001 were considered significant.

### Proteome analysis

Unbiased analysis of the cardiac proteome was performed as detailed elsewhere^119^. Briefly, three hearts per experimental group were homogenized into ice-cold isolation buffer (300 mM KCl, 30 mM PIPES pH 6.6, 0.5% NP-40, 1× protease inhibitor [Roche], 1× phos-stop [Roche]), followed by centrifugation at 15,000 rpm for 10 minutes in a tabletop centrifuge at 4°C to remove insoluble proteins. Supernatants were diluted 1:4 with ice-cold dilution buffer (1× phos-stop [Roche], 0.5% NP-40, 1 mM DTT) and centrifuged again at 15,000 rpm for 15 minutes at 4°C to precipitate actomyosin components. The supernatants were snap frozen for subsequent mass-spectrometry and immunoblot analyses.

Identification of proteins by mass-spectrometry was done by diluting of proteins into TNE buffer (50 mM Tris-HCl pH 8.0, 100 mM NaCl, 1 mM EDTA) and addition of RapiGest SF reagent (Waters Corp.) to a final concentration of 0.1% before boiling of samples for 5 minutes. Following denaturing of proteins, TCEP (Tris (2-carboxyethyl) phosphine) was added to a final concentration of 1 mM and samples were incubated at 37°C for 30 minutes. Samples were carboxymethylated with 0.5 mg/ml of iodoacetamide for 30 minutes at 37°C followed by neutralization with 2 mM TCEP (final concentration). Proteins were digested with trypsin (trypsin/protein ratio of 1:50) overnight at 37°C. RapiGest was removed by adding 250 mM HCl at 37°C for 1 hour followed by centrifugation at 18,000 g for 30 minutes at 4°C. Soluble peptides were extracted and desalted using C18 desalting strips (Thermo Scientific). Trypsinized samples were labeled with sample-specific isobaric tags (iTRAQ8, ABSCIEX)^120^. Each set of smaples was pooled and fractionated using high pH reverse-phase chromatography (HPRP-Xterra C18 reverse phase, 4.6 mm × 10 mm, 5-μm particle, Waters) using chromatography conditions described elsewhere^119^. Subsequently, pooled fractions were analyzed by high-pressure liquid chromatography coupled with tandem mass spectroscopy (LC-MS/MS) using the nanospray ionization method on a TripleTof 5600 hybrid mass spectrometer (ABSCIEX) interfaced with nanoscale reversed-phase UPLC (Waters Corp. nano ACQUITY) equipped with a 20-cm 75-μm ID glass capillary containing 2.5 μm C18 (130 Å) CSHTM beads (Waters Corp.). Peptides were eluted from the C18 column into the mass spectrometer using a linear gradient (5%–80%) of acetonitrile at a flow rate of 250 μl/min for 1 hour. MS/MS data were acquired in a data-dependent manner in which the MS1 data were acquired for 250 ms at m/z of 400–1250 Da, and the MS/MS data were acquired from m/z of 50–2,000 Da. Data acquisition parameters were: MS1-TOF acquisition time of 250 ms, followed by 50 MS2 events of 48-ms acquisition time for each event. The threshold to trigger a MS2 event was set to 150 counts when the ion had the charge state +2, +3, and +4. The ion exclusion time was set to 4 seconds. The collision energy was set to iTRAQ experiment setting. Finally, the collected data were analyzed using Protein Pilot 5.0 (Absciex) for peptide identifications and Peaks Studio (version 8.5, Bioinformatics Solutions Inc.)^121^. Observed changes to protein amounts with a Significance of > 5 and no missing values for identified proteins were considered significant.

### Immunoblot analyses

For analysis of protein expression levels in total cardiac extracts, hearts were homogenized directly into a modified version of SDS-sample buffer^106^. Samples were run on Tris-Glycine SDS-PAGE gels and transferred by electrophoresis onto nitrocellulose membranes (Bio Rad). Following transfer, membranes were stained with Ponceau solution (Sigma-Aldrich) to visualize proteins. Ponceau stained membranes were imaged. Membranes were blocked with 5% BSA dissolved in low salt (0.9% NaCl, 10mM Tris-HCl pH7.4, 0.2% Tween-20) for 2 hours at room temperature with gentle agitation. Membranes were incubated with primary antibody diluted into blocking solution over night at 4°C on a shaking platform. Following the incubation, membranes were washed three times in low salt solution before incubation with secondary antibody diluted in blocking solution for 1 hour at room temperature on a shaking platform.

Subsequently, membranes were washed five times for 10 minutes each with low salt, before being developed using Super Signal West Pico Plus solution (Thermo Fisher Scientific) and imaged on a ChemiDoc Gel Imaging system (Bio Rad). Densitometry on bands was performed using Bio Rad Image Lab (version 5.2.1, Bio Rad) or ImageJ (NIH, version 1.53).

### Echocardiography, hemodynamics and pressure-volume loop analysis

Cardiac functions were measured by transthoracic echocardiography using a Fujifilm VisualSonics Vevo 2100 ultrasound system with a 32-55-MHz linear transducer. Semiconscious anesthetized mice are imaged in M-mode, 2-dimensional, and Doppler echocardiography, which allows for measurements of chamber dimensions, wall thicknesses, mitral inflow and tissue Doppler parameters. Small electrodes were inserted into one upper and one lower limb, which allowed for simultaneous recording of electrocardiograms. Fractional shortening was calculated as an indicator for systolic cardiac function. Heart rate, interventricular septal thickness during diastole (IVSDd), left ventricular posterior wall thickness during diastole (LVPWd), and left ventricular internal dimensions during diastole and systole (LVIDd and LVIDs, respectively) were recorded and analyzed as previously described ^119, 122^.

The procedures to conduct invasive hemodynamics and pressure-volume (P/V-loop) analyses were previously described^123^. After bilateral vagotomy, a 1.4F micromanometer catheter (Millar Inc) was inserted retrograde into the aorta via the right carotid artery and advanced into the left ventricle of anesthetized and intubated mice. Administration of graded dobutamine doses of 0.75, 2, 4, 6, and 8 µg/kg/min is delivered infusion pump (PHD 2000, Harvard Apparatus) for 3 minutes at each dose through the cannulated femoral vein. Baseline pressure measurements are obtained. Data are reported after bilateral vagotomy. Left ventricular hemodynamic parameters are recorded and analyzed using LabChart (ADInstruments, Inc.), including peak left ventricular pressure, left ventricular end-diastolic pressure (EDP), left ventricular dP/dt max (an index of systolic function) and dP/dt min (an index for diastolic distensibility) and Tau (time constant of left ventricular relaxation).

For P/V-loop analyses, baseline echocardiograms were recorded of anesthetized and intubated mice, before administration of 2mg/ml of Ivabadrine-HCl (1.7mg/kg body weight) through the femoral vein. A conductance catheter-micromanometer (Millar Instruments) was inserted into the left ventricle through the inferior vena cava. The P/V catheter was calibrated using echocardiography. During P/V-loop recording, stepwise bolus infusion of saline into the animals blood circulation with volume steps of 250 µl, 375 µl, 500 µl, 750 µl, and 1000 µl was performed through the femoral vein, with an additional echocardiographic recording done after infusing a total of 750µl saline. The circulation was left to recover after each bolus of saline infusion for at least 5 minutes. Following bolus infusion, the inferior vena cava was briefly occluded three times. Following the removal of the catheter, an additional echocardiographic reading was performed. Analysis of data was performed as previously described^123, 124^.

### Hydroxyproline Assay

Total hydroxyproline content to quantify extracellular matrix deposited collagen was measured using a commercially available kit (#MAK008, Sigma-Aldrich) as previously detailed^125^. In short, PFA-fixed cardiac tissue was homogenized in water (10µl water/mg tissue). The lysate was incubated with an equal volume of 12M HCl at 120°C for 3h. 20-50µL of the cardiac homogenates were dried before incubation with chloramine-T and p-dimethyl-amino-benzaldehyde at 60°C for 90 min. Absorbance was measured at 560 nm in a colorimetric assay using a SpectraMax photospectrometer (Molecular Devices) and collagen content of samples was determined using a standard curve generated with hydroxyproline provided in the kit.

### Antibodies

The list of primary antibodies used in this study can be found in **Supplemental Table 1**. Secondary antibodies for immunofluorescence or horseradish peroxidase-linked secondary antibodies for immunoblot detection were either from Jackson Immuno Research Labs or Cell Signaling.

### Serial block face scanning electron microscopy (EM)

The method to prepare and image serial block face electron micrographs has been described in detail elsewhere^126^. Hearts from 9 months old male animals from each group were perfused with Ringers Solution containing xylocaine (0.2mg/ml) and heparin (20 units/ml) for 2 minutes at 35°C, before being fixed by perfusion using 0.15M cacodylate buffer (pH 7.4) containing 2% formaldehyde and 2.5% glutaraldehyde (Electron Microscopy Sciences) as well as 2mM CaCl_2_ at 35°C for 5 minutes. Hearts were then removed and fixed for an additional 2-3 hours on ice in fixative solution. Further processing for electron microscopy by using solutions containing osmium tetroxide, thiocarbohydrazide, uranyl acetate, and lead aspartate was done as described by Wanner, Genoud and Friedrich^127^. Samples were dehydrated in ethanol and transferred through several washes with acetone to be finally embedded in Durcupan resin (Electron Microscopy Sciences). Imaging was done using a serial block face scanning electron microscope by Zeiss with a step-size of 0.1µm, using the 3View system (Gatan). Image stacks of at least 4 cells were imaged per sample. The image stacks were processed for reconstruction and analysis of sarcoplasmic reticulum and mitochondrial volumes using Imaris (Oxford Instruments).

### Atg4 in vitro peptidase assay

For analysis of Atg4 peptidase kinetics, GST-tagged versions of Atg4b, Obsl1 Ig-domains 19-20, Lc3, Gabarap-L1 and GST itself were expressed in freshly transformed E. coli BL21 cells (C601003; Invitrogen) as described in Blondelle et al.^128^ Following induction with 0.2mM IPTG and overnight expression at 18°C, cells were pelleted and lysed into lysis buffer (150 mM NaCl, 10 mM Tris–HCl pH 8, 1% Triton X100). Following lysis, cells were sonicated for 1 minute at 50% output on ice using a Vibracell Sonicator (Sonics & Materials Inc.). Lysate was centrifugated for 45 minutes at 11,000 rpm in an ultracentrifuge (Sorvall) to pellet insoluble proteins. The supernatant was incubated with 200µl Glutathione-Sepharose beads (bioPLUS fine research chemicals) for 2h at 4 °C with agitation. After washing of beads with ice-cold 1xPBS for three times, bound protein was eluted with GST elution buffer (150 mM NaCl, 50 mM Tris–HCl pH 7.4, 150 mM reduced glutathione) and dialyzed overnight at 4 °C against dialysis buffer (150mM NaCl, 20mM Tris-HCl pH8, 1mM DTT, 2mM CaCl_2_). Concentration of dialyzed proteins was determined using a standard Bradford colorimetric assay (Bio Rad) according to the manufacturer’s instructions, and protein concentrations were normalized by diluting proteins in dialysis buffer.

For in vitro peptidase assays, protein combinations were diluted in reaction buffer (50 mM NaCl, 20 mM Tris-HCl pH8, 1 mM DTT, 2 mM CaCl_2_) on ice. Proteins were incubated for 30 minutes at 37°C before addition of an equal volume of modified SDS-sample buffer^106^. Samples were run on SDS-PAGE and proteins were visualized using Coomassie stain. Stained proteins were imaged on a flatbed scanner and quantified by densitometry using ImageJ (NIH, version 1.53).

### Bioinformatics analyses, visualization of data and statistics

Analysis of RNA-Seq data was done in the R environment (v. 4.1.0, https://www.R-project.org/) with packages for MultiQC^115^, STAR^116^, featureCounts^117^ and DESeq2^118^. Analysis of proteome data was done using Peaks Studio (version 8.5, Bioinformatics Solutions Inc.)^121^ also using the built-in statistics. Enrichment analysis and hierarchical clustering were done by using Metascape (http://metascape.org)^129^ and Morpheus (https://software. broadinstitute.org/Morpheus), respectively. Volcano plots, bar graphs and histograms were assembled using Prism (version 9; GraphPad Software) and Excel (Microsoft). If not stated otherwise, data are presented as mean values with standard error of mean (SEM), and statistically significant differences between groups were assessed using either T-test or analysis of variance (ANOVA), as appropriate. P-values of < 0.05 were considered as statistically significant. Sample sizes are detailed either in the figure or figure legend. If not state otherwise, both sexes were analyzed in the experimental procedures.

## Supporting information

Supplemental File 1 - RNA-Seq data

Supplemental File 2 - Matlab script

Supplemental File 3 - Proteome data

Supplemental File 4 - Proteome comparison

## Data availability

All key data supporting the conclusion are presented in main figures, table, or Supplemental Material. Obscurin and Obsl1 gene targeted mice used in this study have been deposited at The Jackson Laboratory, with research identifiers IMSR_JAX:035303 and IMSR_JAX:035302, respectively. Raw next generation sequencing and proteome datasets as well as unmodified original immunoblots and other relevant datasets are available from the corresponding author upon request.

## Acknowledgements

We like to express our gratitude to Alexander Lun, Dr. Stephanie Myers, Anush Velmurugan, Matthew Wright, Antonio Kourieh, Agnieszka Brzozowska-Prechtl and Dr. Jennifer Santini for technical assistance, to Dr. Mathias Gautel for provision of the titin M8 antibody and to Dr. Sorrentino for provision of the sAnk1.5 antibody.

Imaging was done at the UC San Diego Microscopy Core Facility and with the help of Dr. Jennifer Santini and Agnieszka Brzozowska-Prechtl.

Sequencing was performed by the SNP&SEQ Technology Platform in Uppsala, Sweden. The facility is part of the National Genomics Infrastructure (NGI) Sweden and Science for Life Laboratory (SciLife Lab). We are grateful to the Developmental Studies Hybridoma Bank, created by NICHD/NIH and maintained at The University of Iowa, Department of Biology, Iowa City for the provision of antibodies.

## Supplemental Figure Legends, supplemental tables and description of supplemental files

**Supplemental Figure S1.**
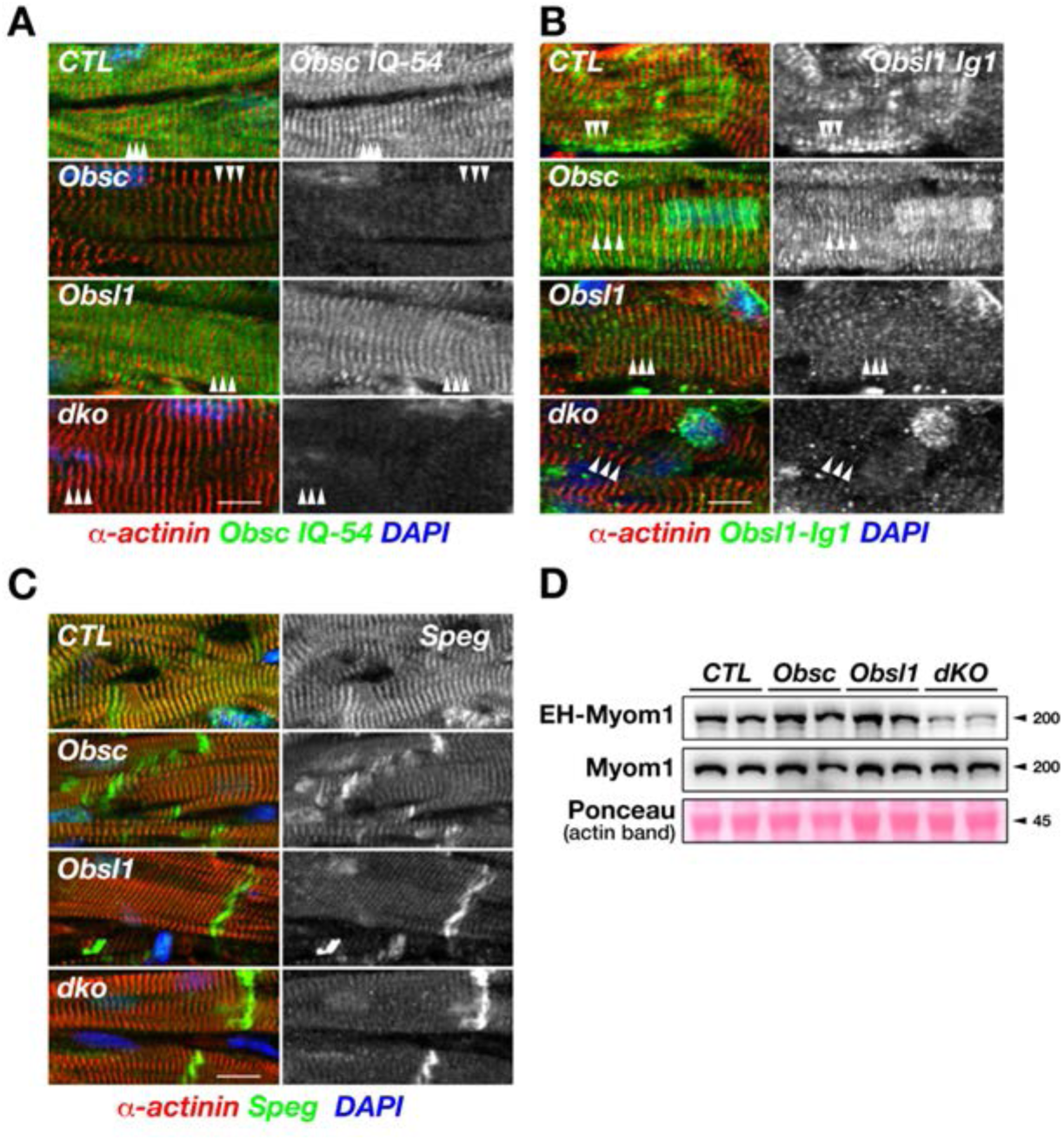
Changes to obscurin, Obsl1 and Speg localization as well as Myom1 expression in single and double knockout (dKO) mice. A-C. Immunofluorescence of frozen cardiac sections from hearts of wildtype (CTL), obscurin knockout (Obsc), Obl1 knockout or obscurin/Obsl1 double knockout (dKO) mice. Sections were stained with antibodies directed against obscurin (Iq-54 epitope, green in the overlay; A), Obsl1 (Ig-domain 1 epitope, green in the overlay; B) or Speg (green in the overlay; C). Dapi (blue) and sarcomeric alpha-actinin-2 were used as counterstains. Scalebar = 10µm. D. Immunoblot analyses of total Myomesin 1 (Myom1) and EH-Myomesin 1 splice isoform protein expression levels in total cardiac extracts of all groups. Ponceau stained actin band is used as loading control.

**Supplemental Figure S2.**
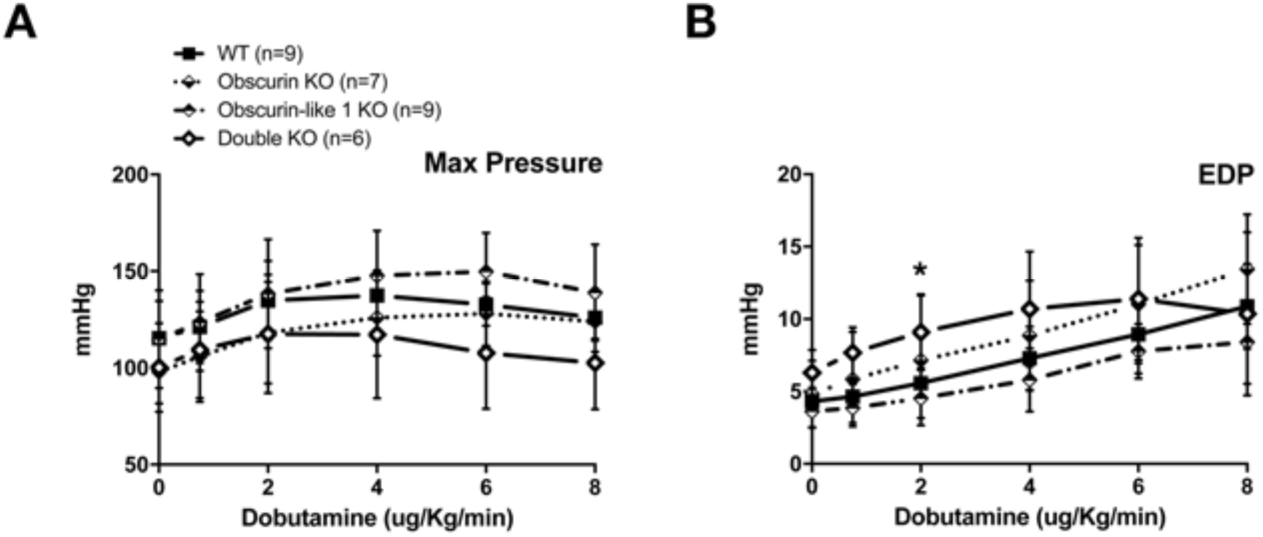
A-B. Hemodynamics analyses of all groups at baseline and with increasing dobutamine amounts. The graphs depict changes to maximum pressure (Max Pressure) and end diastolic pressures (EDP). Shown are averages with standard error of mean (SEM) values. * p < 0.05 vs CTL by two-way ANOVA analyses, with multiple comparisons using Dunnett’s test.

**Supplemental Figure S3.**
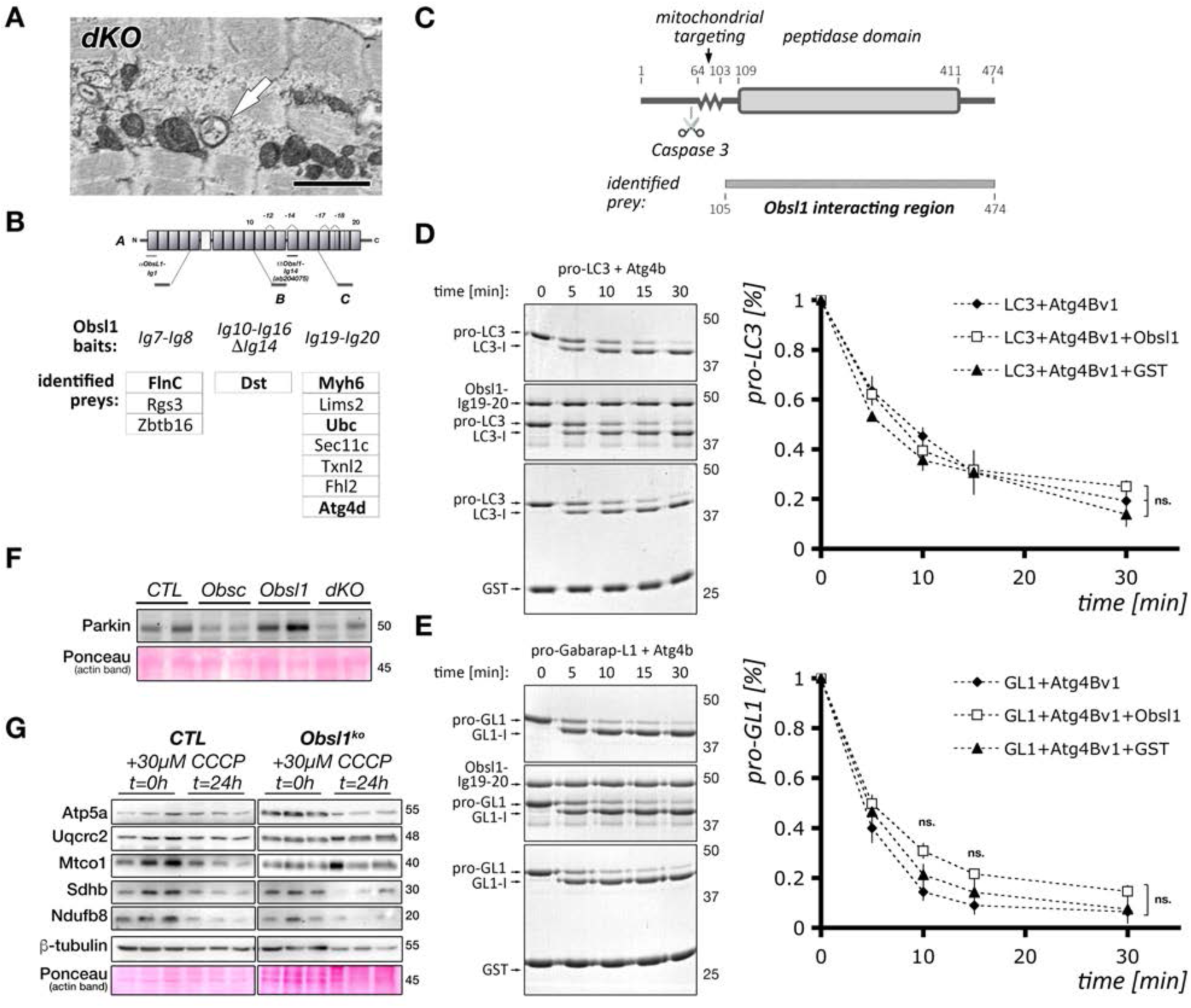
Obsl1 modulates cellular mitophagy through Atg4d. A. EM micrograph of obscurin/Obsl1 dKO heart tissue. Scalebar = 2µm. Arrow indicates double membraned autophagosome. B-C. Schematic overview of Obsl1 interaction partners identified through yeast two-hybrid screens (B; adapted from^43^) and minimal binding site of Obsl1 Ig19-20 domain interacting motif in Atg4d (C). Domains and motif borders/sites in mouse Atg4d delineated by amino acid residues. D-E. Obsl1 Ig19-20 binding to Atg4b does not influence peptidase function for processing of pro-Lc3 (D) or pro-Gabarap-L1 (GL1; E) to the mature Lc3-I or GL1-I forms of the proteins, respectively). Representative Coomassie stained gels (left panels in D, E) and quantitative analysis of average pro-Lc3 or pro-GL1 levels (in % of t=0 minutes; right panels in D, E) with standard errors of mean are shown. Sample size = 3 independent experiments. Statistical significance between groups was tested with ANOVA, with multiple comparisons using Dunnett’s test for each timepoint. F. Immunoblot analysis of Parkin protein levels in cardiac extracts of wildtype (CTL), obscurin or Obsl1 knockout and obscurin/Obsl1 double knockout (dKO) mice. Ponceau stained actin serves as a loading control. G. Representative immunoblots assessing changes to electron transport chain and structural mitochondrial proteins by 30µM CCCP in Obsl1 knockout and control lung endothelial cells at t=0h and t=24h. Tubulin and Ponceau stained actin bands are shown as loading controls.

**Supplemental Figure S4.**
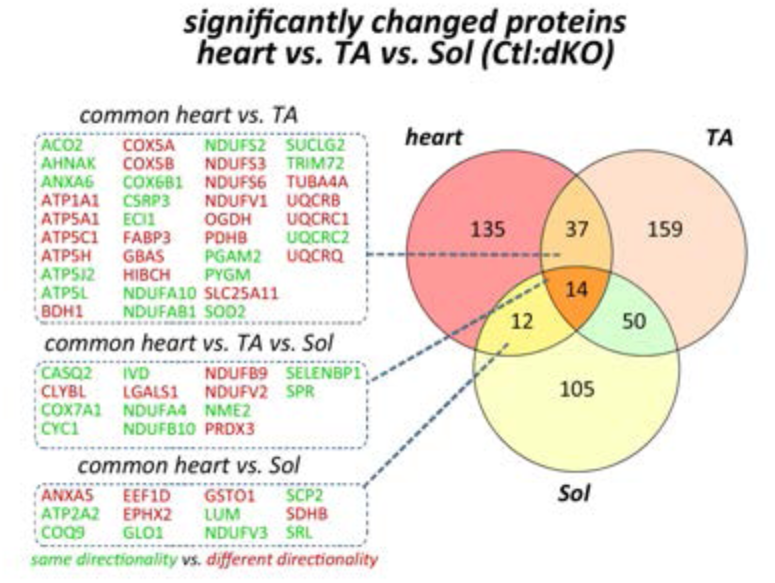
Comparison of significantly deregulated proteins (compared to wildtype controls) using proteome data of cardiac as well as skeletal muscle-specific obscurin/Obsl1 dKO mice. For skeletal muscle-specific dKO, soleus and tibialis anterior muscles were analyzed. This Figure is related to Supplemental File 4.

**Supplemental Table 1.** Primary antibodies used in this study.

| Target | Source and comments |
| --- | --- |
| Obscurin | Rabbit polyclonal antibody with an epitope directed against the region of the Iq-motif to Ig54 of obscurin. Generated in-house and knockout validated<sup>23</sup>. |
| Obsl1 | Rabbit polyclonal antibody with an epitope directed against Ig-domain 1 of human Obsl1. Generated in-house and knockout validated<sup>43</sup>.<br>ab204075; Abcam; epitope targeting aa 1267-1355 (Ig-domain 14); knockout validated<sup>43</sup> |
| Speg | HPA018904, Sigma-Aldrich<br>sc-167091, Santa Cruz Biotechnology |
| Titin M8 | kind gift of Dr. Mathias Gautel; King's College London |
| Sarcomeric alpha-actinin 2 | clone EA-53; Mob 227-05; Diagnostic Biosystems |
| Collagen-I | ab34710, Abcam |
| Smooth muscle-Actin | clone 1A4, DAKO |
| sAnk1.5 | Rabbit polyclonal antibody generated in the Sorrentino laboratory; knockout validated<sup>90,130</sup><br>ARP42566_T100; Aviva Sysbio |
| Sarcalumenin | clone XIIC4; Developmental Studies Hybridoma Bank; developed by K. P. Campbell |
| Histidine-Rich Calcium Binding Protein | SAB1303636, Sigma Aldrich<br>sc-138023, Santa Cruz Biotechnology |
| Calsequestrin 1/2 | ab3517, Abcam |
| Phospholamban (PLN)<br>total- and phospho-PLN | Total phospholamban: MA3-922, Thermo Scientific<br>PLN-S16: A010-12AP, Badrilla<br>PLN-T17: sc-17024-R, Santa Cruz Biotechnology |
| RyR2 | ARR-002, Alome Labs |
| DHPR | ab2864, Abcam, knockout validated |
| Hexokinase (Hk) | Hexokinase I: #2024, Cell Signaling<br>Hexokinase II: #2867, Cell Signaling |
| Phosphofructokinase (Pfk) | • #8164, Cell Signaling |
| Pyruvate kinase M2 (Pkm2) | • #4053, Cell Signaling |
| Phospho- and total Pyruvate Dehydrogenase (Pdh) | • Total Pdh: #3205, Cell Signaling<br>• Phospho-Pdh Ser 293: #31866, Cell Signaling |
| Bdh1 | • 15417-1-AP, Proteintech |
| Scp2 | • 23006-1-AP, Proteintech |
| Acadm | • 55210-1-AP, Proteintech |
| Acetyl-Lysine | • #9441, Cell Signaling |
| Mitofusins 1 & 2 | • Mitofusin 1: #14739, Cell Signaling<br>• Mitofusin 2: #11925, Cell Signaling |
| Phospho-Drp1 and total Drp1 | • Total Drp1: #8570, Cell Signaling<br>• Phospho-Drp1 Ser 616: #4494, Cell Signaling |
| Chchd3 | • OAAN03902, Aviva Sysbio |
| Mic13 | • 25514-1-AP, Proteintech |
| Mitofilin (Mic 60) | • 10179-1-AP, Proteintech<br>• ab110329, Abcam |
| Sam 50 | • WH0025813M4, Sigma-Aldrich<br>• 28679-1-AP, Proteintech |
| Oxphpos antibody cocktail | • 45-8099; Novex; Life Technologies |
| Mitochondrial biogenesis antibody cocktail | • ab123545, Abcam |
| Cox 4 | • #4844, Cell Signaling |
| Lc3a | • #4599, Cell Signaling |
| Atg4b | • ab154843, Abcam; knockout validated<br>• #5299, Cell Signaling; RNAi validated |
| Atg4d | • ab237751, Abcam<br>• NBP3-03374, Novus Biologicals |
| Pink1 | • #6946, Cell Signaling |
| Parkin | • #4211, Cell Signaling |
| Nix/Bnip3L | • #12396, Cell Signaling |
| P62 | • ab155686, Abcam; RNAi validated<br>• ab56416, Abcam; Knockout validated<br>• 610832, BD Transduction Laboratories |
| Phospho and total Ire1 alpha | • NBP2-50067, Novus Biologicals |
| Gm130 | • 610822, BD Transduction Laboratories |
| Xbp1 | • ab220783, Abcam |
| Myomesin 1 (Myom1), EH-myomesin | • Mouse monoclonal Myomesin B4, generated in the lab of Prof. Jean-Claude Perriard, ETH Zurich<sup>131</sup>; kind gift of Dr. Elisabeth Ehler or |
|  | Developmental Studies Hybridoma Bank, developed by Perriard, J.-C.<br>• Rabbit polyclonal EH-Myomesin, generated by Dr. Irina Agarkova in the lab of Prof. Jean-Claude Perriard, ETH Zurich<sup>132</sup> |
| Flag epitope | • F3165, Sigma-Aldrich |
| Beta-Tubulin | • clone E7; Developmental Studies Hybridoma Bank; developed by Klymkowsky, M. |

### Supplemental File 1

Analysis of RNA-Seq data from hearts of wildtype (CTL), obscurin (Obsc), Obsl1 and obscurin/Obsl1 double knockout mice (dKO). Three biological samples from 3-6 months old male and female mice were tested per group.

### Supplemental File 2

Custom Matlab script to analyze calcium transient traces in neonatal mouse cardiomyocytes.

### Supplemental File 3

Proteome analysis of hearts from wildtype (CTL), obscurin (Obsc), Obsl1 and obscurin/Obsl1 double knockout mice (dKO). Three pooled biological samples from 6 months old male mice were tested per group.

### Supplemental File 4

Comparison of significantly deregulated proteins identified by proteome analysis in heart, soleus and tibialis anterior muscles of cardiac- and skeletal-muscle specific dKO mice, respectively.

