## Supplemental File 2 - Matlab script for "Combined loss of obscurin and obscurin-like 1 in murine hearts results in diastolic dysfunction, altered metabolism and deregulated mitophagy": CA_TRANSIENT.m

```
%CA_TRANSIENT
%  Analyzes fluo-4 line-scan traces
%  Values must be imported as numeric matrix (not table) with data in columns
%  times in [s]
%  values in raw fluorescence intensity values
%
%  example:
%
%  (time1)  (value1)  (time2)  (value2) ... headers do not get imported
%  0        250.23     0        1500.35
%  0.001    253.21     0.0021   1503.21
%  .        .          .        .
%  .        .          .        .
%
%
% Usage:
% [n , np, ptp, a, w, rv, ttp, tau, ninetyp]=CA_TRANSIENT(save_tofile, trace_matrix, smooth_value, minpeak_value);
%
% n   - line scan no.
% np  - number of detected peaks & position (note: first and last peak will
%       not be used for calculations)
% ptp - peak-to-peak values, averages & StdErrors (ms), 
%       'variability'; values closer to 0 indicate more 'variability'
%       determined by calculating square of -log(ptp.stddev divided by
%       ptp.average); modified from Berger et al. Circulation 1997
%       doi: 10.1161/01.CIR.96.5.1557
% a   - amplitudes for peaks, average amplitudes & StdError [AU(F/F0)]
% w   - width at half-amplitude
% ttp - time-to-peak values, average TtP & StdError [ms]
% rv  - max rise velocities, average rv & StdError [deltaAU(F/F0)/ms]
% tau - tau decay values, average tau(decay) & StdError [ms]
% ninetyp - times for 90% return to baseline, average & StdError [ms]
%
% Parameters
%
% save_tofile       - save graphs & data to file (0=no; 1=yes)
%                     files will be saved in folder 'results' in working
%                     directory of MATLAB;
%                     data (values only) are saved in xls format (or cvs)
%                     with individual measurements in rows for each
%                     line-scan; graphs are saved as png;
% trace_matrix      - data matrix (see above)
% smooth_value      - lowpass filter value for smoothing (e.g.: 10)
% minpeak_value     - cutoff for peak detection (e.g. 0.5)

function [n , np, ptp, a, w, rv, ttp, tau, ninetyp]=CA_TRANSIENT(save_tofile, trace_matrix, smooth_value, minpeak_value)
%Default settings if no additional parameter provided
%Default settings
%p.smooth_value = 10; p.minpeak_value = 0.5;


%if isempty(smooth_value)=='true'; 
% smooth_value = 10; end
%if isempty(minpeak_value)=='true';
%minpeak_value = 0.5; end

%get number of data points and number of measurements
[no.points,no.n]=size(trace_matrix);

n=[no.n./2];  %sets the total number of line scans

if save_tofile==1; 
    mkdir('results');
end;

for r=1:n;    %loop start for each line-scan
    
clearvars intensity intensity_f intensity_ff0 globalF0 pv loc pw pp baseline;

value.time=trace_matrix(2,(r*2)-1)*1000;                      %determine the time increase point-to-point (in ms) from row 2 

for s=1:no.points;
    intensity(1,s)=trace_matrix(s,r*2); %convert data in column to data in row (needed for ct_filter function)
end

%smooth curve, determine global F/F0 values, set time & find peaks
intensity_f=ct_filter(intensity,'low',smooth_value,0);  % lowpass filter to smooth curve (remove high freq. noise)
globalF0=min(intensity_f);                              % get the global F0
intensity_ff0=intensity_f./globalF0;                    % calculate F/F0 for the whole curve
[pv, loc, pw, pp]=findpeaks(intensity_ff0,'minpeakprominence',minpeak_value,'annotate','extents');
np.no(r)=[length(loc)-2];                                  % sets the detected number of peaks, exclude first/last peak
for s=2:length(loc)-1;
    np.time(r,s-1)=[trace_matrix(loc(s),(r*2)-1)*1000];              % sets the time position of individual peaks, exclude first/last peak
end;         
    
%~~~~~~~~~~~~~~~~~~~~~~~~~~~~~~~~~~~~~~~~~~~~~~~~~~~~~~~~~~~~~~~~~~~~~~~~~~~~~~~~~~~
% determine peak-to-peak, average peak-to-peak and stderror, exclude first/last peak
% determine 'variability index'
%~~~~~~~~~~~~~~~~~~~~~~~~~~~~~~~~~~~~~~~~~~~~~~~~~~~~~~~~~~~~~~~~~~~~~~~~~~~~~~~~~~~
clearvars ptp_value;   %clear variable
for s=2:length(loc)-1;
    ptp_value(s-1)=[(loc(1,s)-loc(1,s-1))*value.time];              %set a frame of 100 data points before peak up to peak
    ptp.value(r,s-1)=[ptp_value(s-1)];                              % sets the individual time-to-peak values (in ms)
end;
ptp.mean(r)=[mean(ptp_value)];                                      % sets the mean time-to-peak value (in ms)
ptp.stderror(r)=[mean(std(ptp_value)./sqrt(length(ptp_value)))];    % sets the mean stderror for the time-to-peak value (in ms)
ptp.variability(r)=[(-log(mean(std(ptp_value))./(ptp.mean(r))))*(-log(mean(std(ptp_value))./(ptp.mean(r))))];
%ptp.variability(r)=[(ptp.mean(r)*(np.no(r)+2))./(value.time*length(intensity_ff0))];  % sets the 'variability' value 
                                                                    % time of peak-to-peak average * number of beats 
                                                                    % divided by total length of time 
                                                                    % the close to 1, the more regular the beats 

%~~~~~~~~~~~~~~~~~~~~~~~~~~~~~~~~~~~~~~~~~~~~~~~~~~~~~~~~~~~~~~~~~~~~~~~~~~~~~~~~
%determine average amplitude & stderror; determine peak width at half prominence; 
%exclude first/last peak
%~~~~~~~~~~~~~~~~~~~~~~~~~~~~~~~~~~~~~~~~~~~~~~~~~~~~~~~~~~~~~~~~~~~~~~~~~~~~~~~~
clearvars amplitude width_value;                                            %clear amplitude & width variable 
for s=2:length(loc)-1;
    frame=intensity_ff0(loc(1,s-1):loc(1,s));
    value.min(s-1)=min(frame);
    value.max(s-1)=intensity_ff0(loc(1,s));
    amplitude(s-1)=value.max(s-1)./value.min(s-1);
    width_value(s-1)=pw(s-1)*value.time;
    a.value(r,s-1)=[amplitude(s-1)];                %sets the individual amplitude values (in AU(F/F0))
    w.value(r,s-1)=[width_value(s-1)];                  %sets the individual half-width values (in (ms))
end;
a.mean(r)=[mean(amplitude)];                        % sets the mean amplitude
a.stderror(r)=[mean(std(amplitude)./sqrt(length(amplitude)))]; % sets the mean stderror for the amplitude
w.mean(r)=[mean(w.value(r,:))];                                % sets the mean half-width (in ms)
w.stderror(r)=[mean(std(w.value(r,:))./sqrt(length(w.value(r,:))))];      % sets the mean stderror for the half-width

%~~~~~~~~~~~~~~~~~~~~~~~~~~~~~~~~~~~~~~~~~~~~~~~~~~~~~~~~~~~~~~~~~~~~~~~~~~~~~~~~
% determine average max rise velocity & stderror, exclude first/last peak
%~~~~~~~~~~~~~~~~~~~~~~~~~~~~~~~~~~~~~~~~~~~~~~~~~~~~~~~~~~~~~~~~~~~~~~~~~~~~~~~~

intensity_diff=diff(intensity_ff0);             % calculate the differential - higher values=steeper increase of fluorescence

%determine the max velocity in a window (100 time-points) before peak; exclude first/last peak
clearvars rv_value;   %clear variable
for s=2:length(loc)-1;
    frame=intensity_diff(loc(1,s)-100:loc(1,s));  %set a frame of 100 data points before peak up to peak
    value.maxrise(s-1)=max(frame);                %get the max value in this frame
    rv_value(s-1)=value.maxrise(s-1)./value.time; %calculate rise-velocity 
    rv.value(r,s-1)=[rv_value(s-1)];                %sets the individual rise-velocity values (in AU(F/F0)/ms)
end;
rv.mean(r)=[mean(rv_value)];                         % sets the mean rise velocity
rv.stderror(r)=[mean(std(rv_value)./sqrt(length(rv_value)))]; % sets the mean stderror for the amplitude

%~~~~~~~~~~~~~~~~~~~~~~~~~~~~~~~~~~~~~~~~~~~~~~~~~~~~~~~~~~~~~~~~~~~~~~~~~~~~~~~~
% determine average time-to-peak and stderror, exclude first/last peak
%~~~~~~~~~~~~~~~~~~~~~~~~~~~~~~~~~~~~~~~~~~~~~~~~~~~~~~~~~~~~~~~~~~~~~~~~~~~~~~~~
clearvars ttp_value;   %clear variable
for s=2:length(loc)-1;
    frame=intensity_ff0(loc(1,s)-100:loc(1,s));     %set a frame of 100 data points before peak up to peak
    [minvalue, minpos]=min(frame);                  %get the min value in this frame just before rise
    baseline(s-1)=minvalue;
    ttp_value(s-1)=(loc(1,s)-(loc(1,s)-100+minpos))*value.time; %calculate rise-velocity 
    ttp.value(r,s-1)=[ttp_value(s-1)];                % sets the individual time-to-peak values (in ms)
end;
ttp.mean(r)=[mean(ttp_value)];                         % sets the mean time-to-peak value (in ms)
ttp.stderror(r)=[mean(std(ttp_value)./sqrt(length(ttp_value)))]; % sets the mean stderror for the time-to-peak value (in ms)

%~~~~~~~~~~~~~~~~~~~~~~~~~~~~~~~~~~~~~~~~~~~~~~~~~~~~~~~~~~~~~~~~~~~~~~~~~~~~~~~~
% determine average tau and stderror, exclude first/last peak
% determine 90% return to baseline value, average & stderror
%~~~~~~~~~~~~~~~~~~~~~~~~~~~~~~~~~~~~~~~~~~~~~~~~~~~~~~~~~~~~~~~~~~~~~~~~~~~~~~~~
clearvars tau_value ninetyp_value;    %clear variable

for s=1:101;                                            %gets the time-values for the frame below (=x-values for fitting) 
    x(s,1)=s*value.time;
end;
for s=2:length(loc)-1;
    frame=intensity_ff0(loc(1,s)+10:loc(1,s)+110);       %set a frame of 100 data points after peak (=y-values for fitting)
    for ss=1:101;                                        %converts the frame from row to column
        y(ss,1)=frame(1,ss);
    end;    
    curve=fit(x,y,'exp1');                               %fits the exponential decay curve
    tau_value(s-1)=-1/curve.b;                           %calculate tau (ms) 
    tau.value(r,s-1)=[tau_value(s-1)];                   %sets the individual tau values (in ms)
    ninetyp_value(s-1)=log(((pv(s))*0.1)./curve.a)./curve.b; % calculates the 90% return to baseline value (in ms)
    ninetyp.value(r,s-1)=[ninetyp_value(s-1)];               %sets the individual 90% return to baseline values (in ms)
end;
tau.mean(r)=[mean(tau_value)];                             % sets the mean time-to-peak value (in ms)
tau.stderror(r)=[mean(std(tau_value)./sqrt(length(tau_value)))]; % sets the mean stderror for the time-to-peak value (in ms)
ninetyp.mean(r)=[mean(ninetyp_value)];                             % sets the mean time-to-peak value (in ms)
ninetyp.stderror(r)=[mean(std(ninetyp_value)./sqrt(length(ninetyp_value)))]; % sets the mean stderror for the time-to-peak value (in ms)

%~~~~~~~~~~~~~~~~~~~~~~~~~~~~~~~~~~~~~~~~~~~~~~~~~~~~~~~~~~~~~~~~~~~~~~~~~~~~~~~~
% plot normalized curve (F/F0) & peaks
%~~~~~~~~~~~~~~~~~~~~~~~~~~~~~~~~~~~~~~~~~~~~~~~~~~~~~~~~~~~~~~~~~~~~~~~~~~~~~~~~

h(r)=figure; 
%subplot(2,1,1); 
plot(trace_matrix(:,(r*2)-1),intensity_ff0(:)); hold on; plot (trace_matrix(loc,(r*2)-1), intensity_ff0(loc),'rv');
%subplot(2,1,2); plot(pp,pw);

filename_char=strcat('results/results_linescan',num2str(r),'.png');
if save_tofile==1;  
 saveas(gcf,filename_char);
end;


% summary bar graph of results
end;
figure;
subplot(3,2,1); bar(a.mean); hold on; errorbar(a.mean, a.stderror, 'k.'); title('Average amplitudes');
subplot(3,2,5); bar(rv.mean); hold on; errorbar(rv.mean, rv.stderror, 'k.'); title('Average rise velocities');
subplot(3,2,3); bar(ttp.mean); hold on; errorbar(ttp.mean, ttp.stderror, 'k.'); title('Average time-to-peak values');
subplot(3,2,2); bar(ptp.mean); hold on; errorbar(ptp.mean, ptp.stderror, 'k.'); title('Average peak-to-peak values');
clear errorbar; subplot(3,2,4); bar(ptp.variability); title('Peak-to-peak variability index');
subplot(3,2,6); bar(tau.mean); hold on; errorbar(tau.mean, tau.stderror, 'k.'); title('Average tau values');


% write summary bar graph and all data to files
filename_char=strcat('results/results_summary.png');
if save_tofile==1;  
 saveas(gcf,filename_char);
 xlswrite('results/amplitudes.xls',a.value);
 xlswrite('results/rise_velocity.xls',rv.value);
 xlswrite('results/time_to_peak.xls',ttp.value);
 xlswrite('results/peak_to_peak.xls',ptp.value);
 xlswrite('results/peak_to_peak_variability.xls',ptp.variability);
 xlswrite('results/width_at_half_height.xls',w.value);
 xlswrite('results/tau.xls',tau.value);
 xlswrite('results/90preturn.xls',ninetyp.value);
 
end;
```
